# The role of space in explaining macroecological patterns of microbial abundance

**DOI:** 10.64898/2026.04.15.718693

**Authors:** Adrián Gutiérrez-Arroyo, Aniello Lampo, José A. Cuesta

## Abstract

Understanding the origin of universal macroecological patterns in microbial communities remains a central open question in ecology. A key observation is that species abundance fluctuations across diverse biomes are well described by a gamma distribution, yet the mechanism responsible for this regularity is debated. Prevailing explanations invoke exogenous stochastic forcing, while endogenous interaction-based approaches—grounded in generalized Lotka–Volterra (gLV) dynamics—have so far failed to reproduce this pattern. Here we show that incorporating spatial structure resolves this discrepancy. In a gLV model with constant migration, local abundance dynamics remain inconsistent with a gamma distribution. However, when the community is embedded in a fragmented landscape as a metacommunity—with species dispersing among patches—aggregating abundances across patches yields distributions that closely match the empirically observed gamma form. We further demonstrate that this result can be reproduced by aggregating independent realizations of the constant-migration gLV model, entailing statistical aggregation rather than a specific biological mechanism as the origin of this pattern. Our findings highlight the fundamental role of spatial structure in shaping microbial macroecology, and suggest that observed abundance distributions may largely reflect the spatial coarse-graining inherent in metagenomic sampling.

## I. INTRODUCTION

Microbial communities are composed of diverse species (taxa), whose relative abundances shape the collective functional properties of the ecosystem. For instance, in human-associated biomes, the composition in terms of taxa abundances is closely linked to host physiology, immunity, and health [1, 2]. Similarly, in soils, the dominance of a subset of species over others strongly influences nutrient cycling, fertility, carbon sequestration, and greenhouse gas emissions, thereby affecting ecosystem multifunctionality and contributing to global environmental change [3–5].

Empirical evidence shows that microbial abundances vary over time and across samples, and understanding the mechanisms that regulate this variation is therefore essential to explain the processes mentioned above. Notably, abundance variation exhibits striking regularities across different biomes, which are commonly studied through the lens of macroecology [6–10], that is, in terms of large-scale statistical distributions. Specifically, Grilli*’*s 2020 seminal work [10] unveiled that fluctuations in species abundances, for several real biomes, are well fitted by a gamma distribution.

The universal character of this pattern suggests the existence of some underlying mechanism driving abundance fluctuations, yet its nature remains debated. Undoubtedly, environmental changes, demographic stochasticity, cross-feeding, competition, mutualism, trophic interactions, and many other ecological forces all contribute to the overall behavior of microbial ecosystems. What remains unclear, though, is which of these forces are actually relevant in shaping the temporal fluctuations in abundances and, consequently, the resulting patterns.

One line of research posits that the emergence of abundance patterns is primarily driven by *exogenous* factors, that is, forces acting externally on the internal dynamics of the community. These include abiotic environmental fluctuations, such as changes in temperature, pH, moisture, salinity, or oxygen levels; pulses and heterogeneity in nutrient availability; and also host-related physiological variation and other kinds of episodic disturbances. This view is reflected in the growing dominance of stochastic population models, which account for environmental fluctuations through a (multiplicative) random noise, and which have gained traction because they can reproduce large-scale statistical regularities using relatively simple assumptions and few parameters [10–17].

Nevertheless, the explicative power of this kind of models is very limited. To begin with, noise terms are included *ad hoc*, with noise always assumed either white or at most exponentially correlated—a convenient but unrealistic choice. Environmental fluctuations are hardly of this form. Not only do they exhibit more complicated time correlations, but often they have built-in periodicities (food ingestion, tides, day-night cycles…) that are seldom accounted for. For this reason, as proposed so far, stochastic models become mere effective theories, good for providing quantitative descriptions of microbial communities but useless to discriminate the actual causes of abundance variability in microbial communities.

A second approach bets on species interactions (a purely *endogenous* factor) in seeking an explanation for the origin of fluctuations in microbial systems. According to this line of thought, the presence of hundreds or thousands of species in these microbial communities creates a dynamic that, although deterministic, can be wildly fluctuating—even chaotic [18–24].

That nonlinear dynamics can generate seemingly random fluctuations in ecological communities is an idea that goes back to Robert May*’*s pioneering discovery of chaos in deterministic population models [25, 26]. Chaotic dynamics have even been proposed, e.g., as a possible explanation of the *‘*plankton paradox*’* [27], whereby many plankton species can coexist on few resources. Although for decades the empirical evidence gave little support to the existence of chaotic dynamics in ecology, a recent revision of the methods employed to detect it, as well as an increase in the quality and availability of data, have revealed that chaos is prevalent, especially in high-dimensional systems [28, 29]. Experiments performed in the early 2000s showed that even microbial communities with few species can exhibit chaotic fluctuations [30].

Fluctuations in highly diverse, deterministic microbial communities have been studied through generalized Lotka-Volterra systems using mean-field theory. This approach aims at deriving an equation for a *‘*focal species*’* that subsumes the interaction with the rest of the community through an effective noise term. The features of this noise must be determined in a self-consistent manner [20, 23, 31]. Thus, contrary to standard stochastic models, this approach does derive the noise term from first principles.

Although the resulting equation superficially resembles Grilli*’*s stochastic logistic model (SLM) [10], the similarity stops there. Whereas noise in the SLM is white, noise in the mean-field equation has—as expected from a real noise term—a nontrivial temporal correlation, which is further coupled to the very dynamic of the abundance through the self-consistent condition. As a consequence, the dynamics are quite different. The SLM reproduces the gamma distribution of abundance fluctuations observed in empirical data [10]. The mean-field equation, however, predicts a distribution of species abundance that looks like a power law with exponent −1 [23]. This result agrees with numerical simulations of the whole dynamical system—even if the latter are performed in a stronger interaction regime [24]—but, as we will show later, it is incompatible with the gamma distribution of abundance fluctuations observed in empirical data. The validity of mean-field theories requires large numbers of species, such as those existing in microbial communities. For this reason, it is noteworthy that the community pursuing this approach had been systematically ignoring the fact that their theory cannot reproduce one of the most ubiquitous and reproducible empirical patterns. To explain the origin of this discrepancy and eventually show how it can be circumvented is the main goal of the present contribution.

Since May*’*s pioneering work [32] it has been known that fluctuating communities undergo species extinctions. In the studies exploring deterministic chaos in these systems, this is prevented by opening them to migration. As a consequence, species alternately thrive or go extinct, in quite an unpredictable manner [23, 24]. This turnover of species maintains a small community of abundant species that respects May*’*s stability limit at all times. As a matter of fact, the log-abundance distribution reflects this dynamics by exhibiting two peaks—at high and low abundances—joined by a low probability plateau—which explains the exponent −1 of the abundance distribution [23].

The need for a migration term suggests changing the point of view from closed to open systems. Ecological communities are usually embedded in a larger environment that continuously supplies new individuals. Although a constant migration is a simple way to account for this openness, a more realistic model should incorporate the spatial distribution of species as a necessary ingredient. In fact, there is ample evidence of the crucial role played by space in shaping microbial communities and driving the emergence of spatially segregated bacterial clusters [33–36]. Consequently, after analyzing the model with migration, we will eventually lift it to a metacommunity model in which species can diffuse between patches [37, 38]. This framework thus represents a step forward in the exploration of the role of space in microbial modeling, a topic that has long remained comparatively underexplored—although it is currently attracting growing attention [20, 39–42].

A central result of our work is that, even though neither the model with constant migration nor isolated patches of the metacommunity exhibit abundance fluctuations comparable to those observed in empirical communities, aggregating species abundances across all patches of the metacommunity recovers the gamma distribution reported by Grilli. This suggests that this pattern might be a consequence of statistical aggregation rather than the fingerprint of an underlying mechanism [43]. This result highlights the importance of the spatial distribution of microbes, not only in sustaining higher diversity, but also in explaining observed macroecological patterns. As we discuss at the end of this article, current experimental protocols of data acquisition favor the aggregation of spatially dispersed microbes. In fact, one of the most relevant conclusions of the present work is that we should devise new experiments that avoid this aggregation if we want to gain insight into the actual microbial dynamics.

## II. OPEN LOTKA-VOLTERRA MODELS

The gLV equations stand as the classical archetype for studying the temporal evolution of species abundances (also referred to as populations). They can be written as

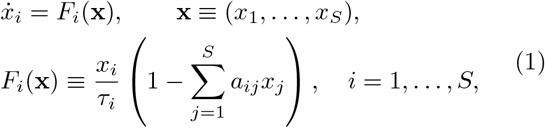

where *x*_*i*_(*t*) is the abundance of species *i* at time *t, τ*_*i*_ is the time scale of its basal population growth, and *S* the number of species *(diversity)* of the community. This model, which represents an isolated ecological community, is sketched in Fig. 1**a**.

**FIG. 1.**
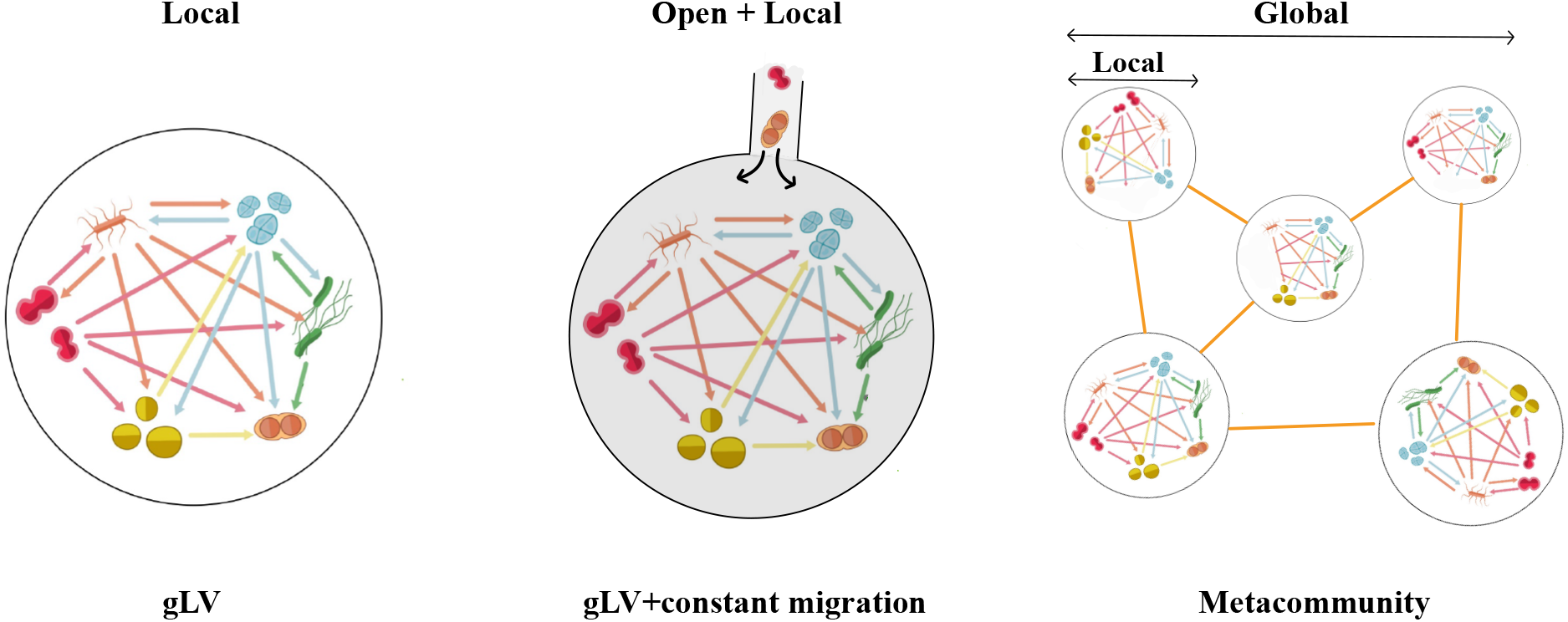
Conceptual illustration of the modeling frameworks. The generalized Lotka–Volterra model neglects spatial structure, describing the community as a well-mixed, fully local system. Introducing constant migration adds an external supply of individuals from the surrounding environment, making the system open but still spatially local. By contrast, the metacommunity framework explicitly incorporates both local and global spatial scales—local patches connected through dispersal within a broader landscape—thus providing a genuinely multi-scale description of ecological dynamics. This conceptual distinction highlights how incorporating spatial structure allows bridging the gap between local population dynamics and largescale macroecological patterns.

The interaction coefficient *a*_*ij*_ represents the effect of the species *j* on the growth rate of species *i*. Matrix *A* = (*a*_*ij*_) can be regarded as the adjacency matrix of the interaction network underlying the microbial community. In this framework, *a*_*ij*_ *<* 0 indicates that species *j* enhances the population growth of species *i*, whereas *a*_*ij*_ *>* 0 inhibits that growth. The different combinations of signs of *a*_*ij*_ and *a*_*ji*_ span all sorts of ecological interactions, from competition (both positive) to mutualism (both negative).

The self-interaction term *a*_*ii*_ *>* 0 is the reciprocal of the carrying capacity for species *i* in isolation, and it can be used to set the scale of species *i’*s abundance. So we will henceforth assume *a*_*ii*_ = 1. As for the time scales *τ*_*i*_ of basal population growth, the model is not very sensitive to variations in these parameters, compared to the effects induced by interactions [40], so we will choose them all equal and scale time with this parameter. Thus *τ*_*i*_ = 1 for all species and time is measured in generations.

Other than the gamma distribution of abundance fluctuations, Grilli reported two other macroecological patterns: a power-law scaling Var(*x*_*i*_) ∝ ⟨*x*_*i*_⟩^2^ across species (Taylor*’*s law) and a lognormal distribution of mean abundances across species ⟨*x*_*i*_⟩. These two patterns are affected by the scaling of the abundances assumed by setting *a*_*ii*_ = 1, however the gamma distribution of abundance fluctuations is blind to it. As understanding the origin of the latter is the primary goal of this paper, we will not concern ourselves with the former (but see the Discussion section V).

In a highly diverse community (*S* ≫ 1) it is not possible to determine empirically the value of the interaction coefficients [44]. To circumvent this problem, Robert May proposed adopting a statistical approach [32], replacing the specific values of the *a*_*ij*_ by randomly generated parameters, so that the mean *µ* and standard deviation *σ* matched those of the actual community. Unless otherwise specified, interspecies interactions *a*_*ij*_ (*i* ≠ *j*) are independently drawn from a normal distribution [45] with given *µ* ≡⟨*a*_*ij*_⟩ and *σ*^2^ ≡Var(*a*_*ij*_). Hence, the overall dynamic of the system will be determined exclusively by the community-level parameters *S, µ*, and *σ*.

It is well established that, in order for this model to exhibit non-trivial fluctuations, the variance of interactions must be sufficiently large [19, 22, 24]. However, both a high number of species and strong interaction couplings render the community unstable [32], leading to species extinctions. Thus, in order to induce a persistent fluctuating dynamic, it is necessary to add a mechanism that precludes extinctions from being definitive. The most natural way to achieve this is to consider open systems. The existence of an environment with which the community interacts allows species that go locally extinct to be restored [20, 23, 24].

By far, the simplest method to emulate an open system is to add a constant migration term to (1) (see Fig. 1**b**):

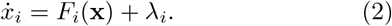

For simplicity, we assume a common migration rate across species, *λ*_*i*_ = *λ* (allowing species-dependent migration rates yields qualitatively similar results). What ultimately matters is primarily the order of magnitude of *λ*, which must remain sufficiently small so as not to supersede the gLV dynamics. In line with the small-migration regime commonly considered in the literature [22–24], we assume *λ* ≤ 10^−4^, ensuring that migration plays only a minimal stabilizing role—preserving large abundance fluctuations while preventing species loss.

A more complex and realistic way to describe an open system assumes that the community is embedded in a fragmented landscape (see Fig. 1**c**) in which *M* identical patches, each containing a replica of the community, are connected forming a spatial network. Species can diffuse from patch to patch through the links. The equations describing such a model are

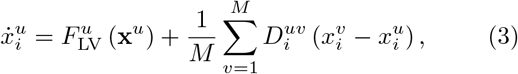

where *i* = 1, …, *S* labels the species, *u, v* = 1, …, *M* label patches, 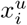 denotes the abundance of species *i* at the patch *u*, and 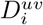 are the rates of migration of species *i* from patch *v* to patch *u*. Matrices 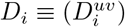 describe the spatial connectivity. This formulation, also known as metacommunity model, describes a system in which species not only interact within each patch according to the gLV dynamics (first term of the right-hand side of (3)), but also disperse across space from patch to patch (second term) [20, 37, 38, 40, 42, 46, 47].

Metacommunities are commonly interpreted as collections of well-defined, isolated patches (such as islands). However, they also arise as discretizations of reaction-diffusion equations describing a continuous spatial distribution of microbes. In this interpretation, patches are simply the points of this discretization. When the characteristic spacing of the discretization is longer than the interaction range of species, its points are effectively isolated from each other. Using this point of view, we can set 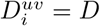 for all species and neighboring patches (and 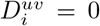 otherwise). Here, we adopt a mean-field approach by assuming that the network of patches is fully connected—thereby discarding any non-trivial landscape topology. Under this simplification the equations adopt the form

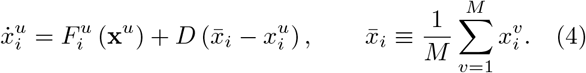

Interestingly, another mean-field argument connects this model with the gLV model with migration, (2). By assuming 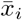 to be constant, we can identify 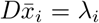 and change the basal growth rate from 1 to 1 − *D*—which can be approximated by 1 because *D* ≪ 1, hence recovering (2).

## III. LOTKA-VOLTERRA WITH MIGRATION

A mean-field analysis of (2) reveals that, depending on the parameters *S* ≫ 1, *µ*, and *σ*, the system can be in one out of three dynamical phases [19]. If—as in the present study—the interaction coefficients are mutually independent, for *σ*^2^*S <* 2 and *µ >* 0 there is a unique stable global equilibrium where *S*^∗^ − *S* species coexist—the remaining *S* − *S*^∗^ go extinct. The fraction *S*^∗^*/S* decreases as *σ*^2^*S* increases up to *σ*^2^*S* = 2, where this fraction becomes 1*/*2—hence 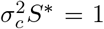, with 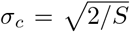 [19, 48]. This is the limit of linear stability of the Lotka-Volterra model, as obtained by May [32]. For *σ*^2^*S >* 2 no stable equilibrium exists anymore and trajectories wander around between fixed points, exhibiting a chaotic behavior. In this regime, abundances fluctuate permanently. Upon further increasing *σ*^2^*S* the system reaches a new threshold value for *σ*^2^*S*—which increases with *µ*. Above that threshold, the abundance of some species diverges in time. Hence, this threshold defines the limit of validity of the Lotka-Volterra model.

This theoretical phase structure found direct support in controlled laboratory experiments with synthetic bacterial communities [22], where community-level parameters alone—species number and interaction strength— were sufficient to predict the emergence of sustained fluctuations under stable external conditions.

For finite values of *S*, there are finite size corrections to this picture. To begin with, these thresholds are no longer abrupt. For *σ*^2^*S >* 2, the probability that a given realization of the interaction matrix (*a*_*ij*_) has a stable fixed point is nonzero [22], although it decreases as we increase either *σ* or *S* (see Fig. 2). In contrast, the observed fluctuating communities can be of two types: periodic or chaotic. At first, the periodic solutions are more likely than the chaotic ones, but eventually (large *σ* or *S*) the latter dominate. Telling them apart can be automated by calculating a Fourier transform of the time series (see *Materials and Methods*).

**FIG. 2.**
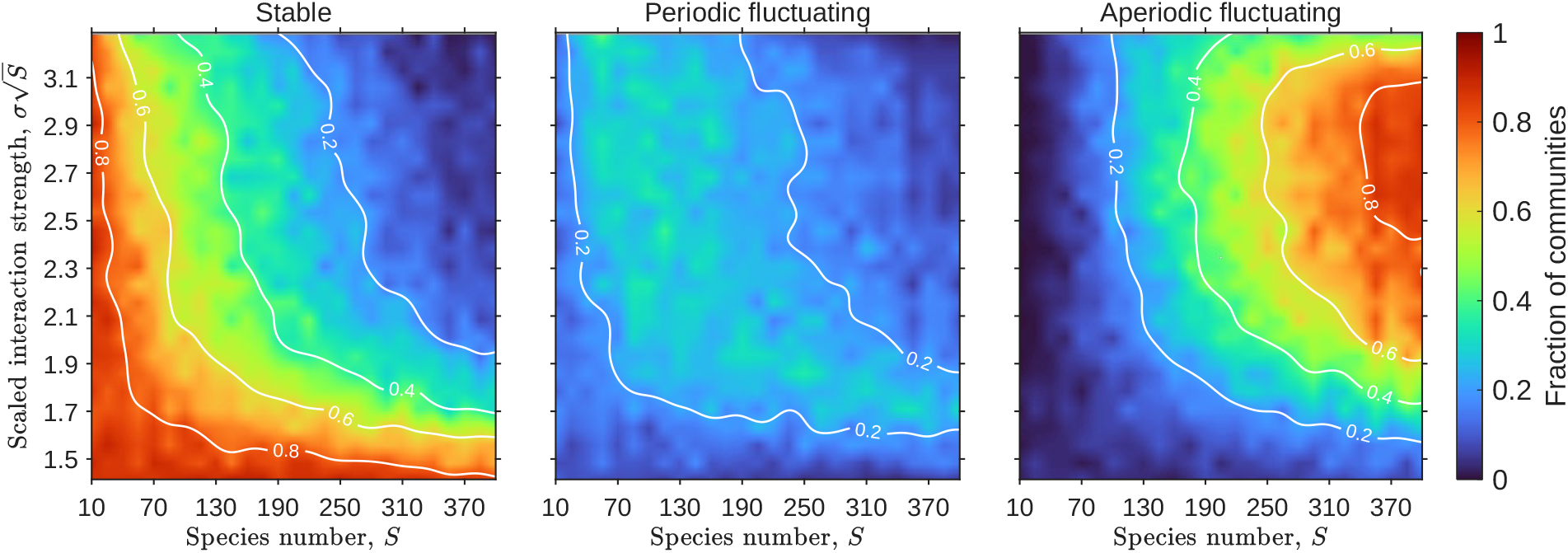
Dynamical phases of the Lotka-Volterra with migration, (2). The number of species *S* and the interaction standard deviation *σ* determine the emergence of stable, periodic, and aperiodic abundance dynamics. The color map represents the fraction of communities displaying the behavior specified in each panel, obtained from ensembles of *N* = 200 simulations and subsequently interpolated for visualization purposes. Trajectories are simulated with parameters *µ* = 10*/S* and *λ* = 10^−8^, and classified as described in *Materials and Methods*. The diagrams show that aperiodic fluctuations become increasingly prevalent as community diversity (*S*) and interaction strength (*σ*) increase. Trajectories that grow unbounded (something that may happen when *σ* is large) are removed from these counts.

In the chaotic regime, the abundance of any species alternates between values *x*_*i*_(*t*) ∼ 1 (the carrying capacity) and values *x*_*i*_(*t*) ∼ *λ* (the migration level). Those with high abundance at a given time *t* form a small community of *S*_eff_(*t*) species such that *σ*^2^*S*_eff_(*t*) *<* 1, thus satisfying May*’*s stability condition [23]. There is a turnover of species in this community, so that its composition changes over time in a seemingly random manner [24] (Figure 3**c** shows the abundance time series of a few species). This turnover causes the distribution of log-abundances *P* (log *x*) to exhibit two peaks, one close to log *x* ≈ 0 and the other close to log *x* ≈ log *λ* (see Figure 3**a**)—although the precise location and height of each peak depend on the interaction parameters (Figs. S2 and S3 of the SI). In between these two peaks the distribution has a plateau—therefore the species abundance distribution (SAD) *P* (*x*) ∼ *x*^−1^ in the interval *λ* ≪ *x* ≪ 1 when *λ* → 0 [23, 24]. Each species has a similar distribution *P* (log *x*) as the whole community, but the locations, and especially the heights of the two peaks, depend on the species, as illustrated by Figure 3**d**. This means that the time spent by different species in the small community varies a lot (Figure 3**e**). Also, the positions of the low (Figure 3**f**) and high (Figure 3**g**) peaks is species-dependent—although they have a smaller variability. We have obtained the results of Figure 3 through numerical integrations of (2), but the solution of the mean-field equations is consistent with them [23, 31].

**FIG. 3.**
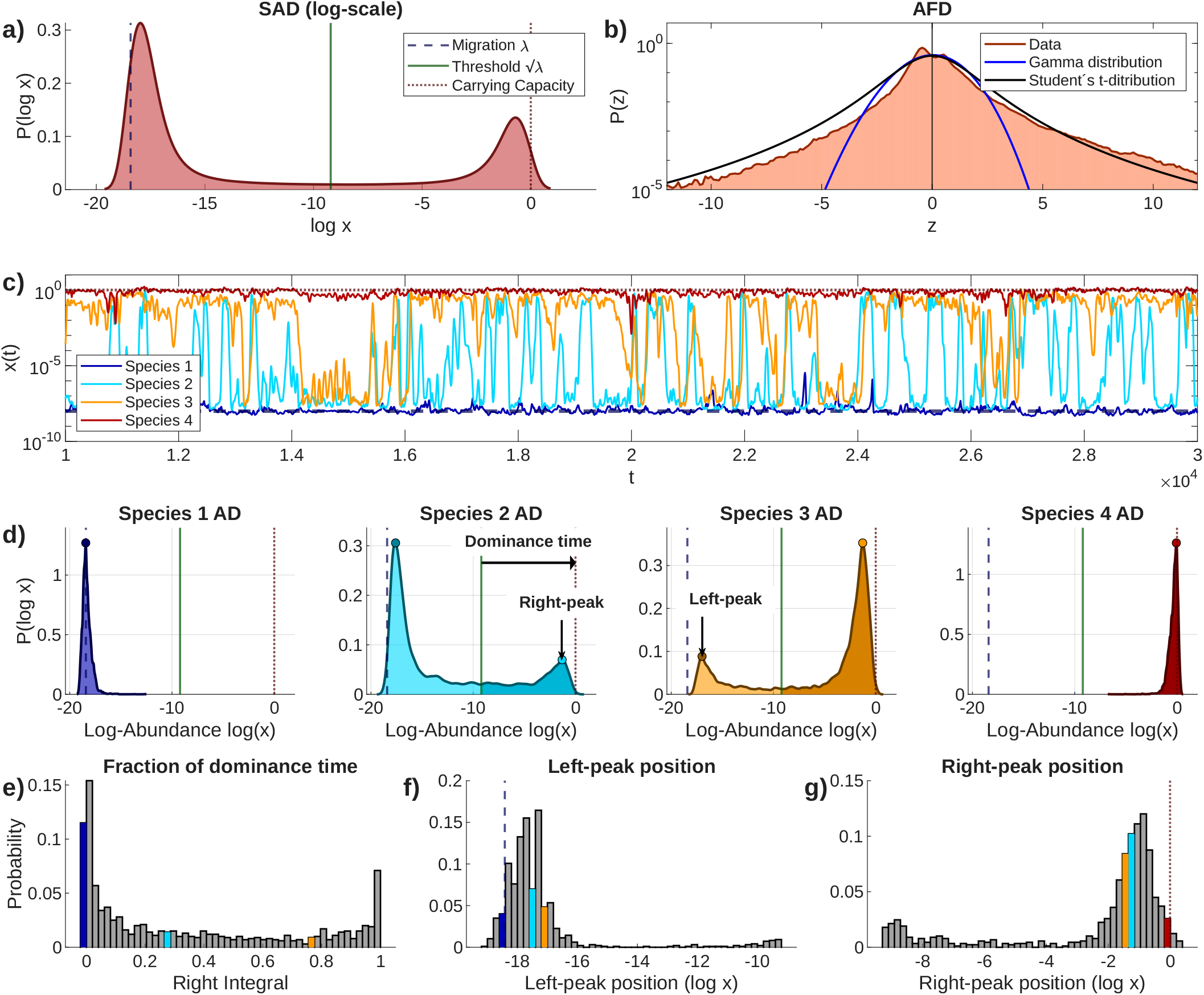
Macroecological characterization of the Lotka-Volterra model with migration, (2). **(a)** Species abundance distribution (SAD) in logarithmic scale, i.e. the distribution of log-abundances across time samples, aggregated over all species in the community. It displays a characteristic bimodal structure with an intermediate plateau. **(b)** Abundance fluctuation distribution (AFD), defined in *Materials and Methods* as the distribution of the standardized log-abundances, (5), aggregated across all species in the community. Solid curves correspond to fits to an exp-gamma distribution, (A2) (blue), and a Student*’*s-*t* distribution (black). The exp-gamma performs very poorly compared to the Student*’*s-*t*. **(c)** Typical abundance time series for 4 selected species. Abundances take values between the migration floor *λ* and the carrying capacity (set to 1), displaying heterogeneous temporal behaviors ranging from small fluctuations to rapid turnover between rare and dominant states. **(d)** Log-abundance distributions for the species shown in **(c)**. These distributions follow the general pattern shown in **(a)**, but the height and location of the peaks varies, reflecting a diversity of behaviors ranging from rare presence to high dominance. **(e)** Histogram of dominance times, defined as the fraction of time during which species abundances lie above 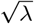 (half the interval in logarithmic scale). **(f), (g)** Distributions of the positions of the left and right peaks of the individual species abundance distributions shown in **(d)**, respectively, and illustrating the heterogeneity of the different species in the community. Insets in **(d)** illustrate the definition of these quantities. Colored bars indicate the locations in t histogram of the species illustrated in **(c), (d)**. Results are obtained from simulations of (2) with *S* = 10^3^, *µ* = 10*/S*, 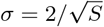, and *λ* = 10^−8^ (the same parameters used in Ref. [23]).

The problem with the phenomenology just described is that the abundance fluctuation distribution (AFD) is not compatible with the distributions obtained from real data. In empirical biomes, the standardized log-abundances

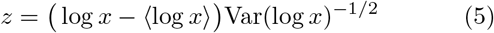

are well described by an exp-gamma distribution (see (A2) in *Materials and Methods*); on the contrary, the distributions obtained from simulations cannot be well fitted by this distribution. Figure 3**b** illustrates how poorly an exp-gamma distribution fits the numerical distribution *P* (*z*); a Student*’*s *t* distribution does a much better job, even though there is no reason to believe it to be the *‘*right distribution*’*. As a matter of fact, for the whole range of parameters *µ* and *σ*, the mean-squared errors of the fits to the logarithm of the numerical AFDs are an order of magnitude larger for the gamma than for the Student*’*s *t* (see Figs. S4 and S5). The reason why a gamma distribution makes a good fit to the empirical data, but not to the simulated time series, is that the former show abundances that fluctuate around some average value, rather than alternating between low and high abundances, as in the latter. And in fact, looking at the distributions of Figure 3**d**, it is clear that the AFD cannot be a gamma. We want to stress that this inconsistency does not arise from the mean-field analysis of this model made by many authors, because the results shown here derive directly from simulations of the Lotka-Volterra system, and they both agree. If we sought a culprit, we should look at the model itself.

Mallmin et al. [24] studied a strong interaction regime, for which *µ, σ*^2^ = *O*(1), rather than *O S*^−1^), as *S* → ∞ [24]. We also studied this regime in our simulations. Our results suggest that the differences between both regimes are quantitative, rather than qualitative. We observe the same turnover of species, which causes the abundance distributions to exhibit two peaks at low and high values, separated by plateaus. The high-abundance community in Ref. [24] is less populated though—hence the high abundance peak is very small (see Fig. S6). However, this is more an effect of the values of the parameters *µ* and *σ* chosen by these authors. Changing them changes the height of the peaks in a continuous manner (Figs. S3 and S7). Needless to say that the AFD is not well described by a gamma distribution in this regime either.

## IV. METACOMMUNITY MODEL

The Lotka-Volterra model with migration is an effective description of a system that exchanges individuals with an external pool. A more realistic approach to an open system amounts to describing our ecosystem as a metacommunity. In this setup, the *‘*external pool*’* is actually made of a set of similar, interconnected communities. Metacommunity Lotka-Volterra models like (4) have also been shown to sustain highly diverse communities in which species abundances undergo persistent aperiodic fluctuations [20, 40, 46, 49]. They also provide a natural way to add space to the modeling of ecosystems. Thus, the model defined by (4) is the obvious next step in our attempt to understand the dynamics of microbial communities.

We simulate (4) with *M* = 30 patches and an initial pool of *S* = 250 species. Choosing *D* needs some consideration. Too high a value of *D* transforms the metacommunity into a well-mixed, isolated community. Therefore, we can expect this system to generate a stable community with few surviving species, in accordance with May*’*s stability limit [20]. On the other hand, too low a *D* renders the patches effectively isolated so that the diversity in each patch is subject to May*’*s limit. We have checked that *D* = 10^−3^ is a value that prevents these two problems and keeps the metacommunity diverse. This notwithstanding, the system tolerates a wide range of diffusion coefficients (see Figs. S11 and S20).

Unlike what happens in an open system with constant migration, species in a metacommunity can go extinct regardless of the value of *D*. Hence, we need to introduce an extinction threshold *x*_ext_. A species is considered extinct, and removed from the metacommunity, once its abundance falls below *x*_ext_ = 10^−20^ in all patches simultaneously; once removed, the species cannot recolonize the metacommunity. This notwithstanding, for all parameter values in the fluctuating regime there is an asymptotic fraction of surviving species 0 *< S*^∗^*/S <* 1 [49] in the metacommunity (see Fig. S8).

The interaction coefficients 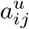 are randomly selected from a multivariate normal distribution such that

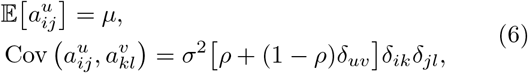

for all *i, j, k, l* = 1, …, *S, u, v* = 1, …, *M*. If *ρ* = 0, coefficients 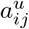 and 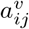, for *u* ?= *v*, are independent; if *ρ* = 1, they are identical. In what follows we choose *ρ* = 0.95, meaning that they have 95% similarity. Justifying this choice requires reflecting on the nature of the interaction coefficients. Generalized Lotka-Volterra models are phenomenological equations aiming at describing an ecosystem in the simplest possible way. Accordingly, the interaction coefficients are nothing but effective parameters that capture the net influence that a species exerts on another one. As such, they encompass not only the species identities, but also all sorts of environmental forces that may affect this interaction. In other words, there is no reason why these effective interactions between a given pair of species should be exactly the same in all patches of a metacommunity. In fact, the choice *ρ* = 1 is not even structurally stable, because the behavior of a metacommunity with *ρ <* 1 (no matter how close to 1) is qualitatively different from another with *ρ* = 1. In the latter, the abundances of a given species in different patches are synchronized—a synchronization that vanishes as soon as *ρ <* 1 [20]. Our choice *ρ* = 0.95 yields *‘*similar*’* interactions between like species across patches, but takes into account local perturbations breaking the symmetry between them.

Figure 4 summarises the results of our simulations. If we look at the results at the *local* scale, i.e. for individual patches, the observed behavior is very similar to what we found for the gLV model with migration. There is the same turnover of species (Fig. 4**b**) and the same two-peaked structure of the distribution of log-abundances *P* (log *x*) (see Fig. S21). Likewise, a gamma distribution is a poor fit to the AFD (Fig. 4**a**). However, the results are quite different when we observe the dynamics of the global (patch-averaged) abundances 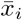 [c.f. Eq. (4)]. To begin with, they no longer exhibit turnover, but fluctuate around an average value (Fig. 4**d**), just like empirical abundances do [10, 12]. Furthermore, the resulting AFD is now well fitted by the distribution given in (A2) (Fig. 4**c**), at least in the range *P* (*z*) ≳ 10^−4^—the resolution threshold of empirical data. Below this threshold, the right tail is well captured by the fit, but the left tail shows a slower decay. We will come back to this point later.

**FIG. 4.**
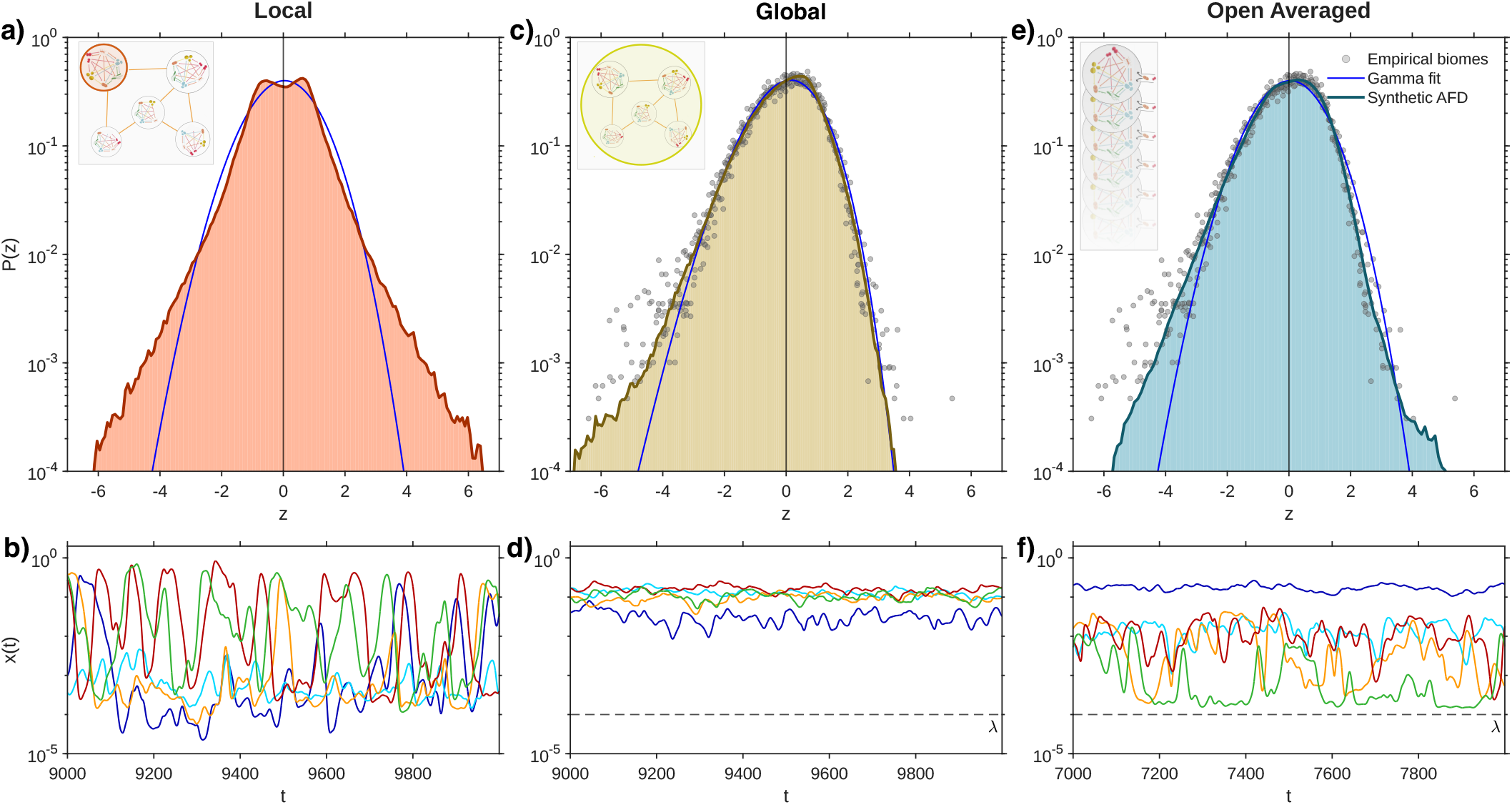
Abundance fluctuation distributions in the metacommunity model, (4). **(a)** Distribution of the rescaled log-abundances, (5), at the local scale, i.e. abundances of species *i* = 1, …, *S* within a single patch 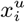. The distribution is very similar to that of a gLV model with constant migration (see Figs. 3 and S4). Clearly, the exp-gamma distribution, (A2), fits poorly the numerical distribution. **(b)** Typical time series of local abundances for selected species, showing the dynamics of species turnover. **(c)** Abundance fluctuation distribution at the global scale in the metacommunity, obtained by averaging species abundances across all patches, 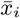 [c.f. Eq. (4)]. Deviations in the left tail aside, the resulting distribution is well described by the exp-gamma, (A2). Grey dots portray empirical biome distributions for *cross-sectional data* selected from the *MGnify* platform [50] (see Fig. S9 for *longitudinal data* and Fig. S10 for a comparison of their fits). **(d)** Corresponding global abundance time series for selected species. In contrast to the local dynamics, these trajectories no longer display turnover but rather fluctuate around their mean value, just like empirical time series. **(e)** Abundance fluctuation distribution obtained by aggregating *M* independent realizations of the constant-migration Lotka-Volterra model, (2), each starting from a randomly drawn initial condition *x*_*i*_ (0) ∼ *U* [0, 1] and with interaction matrices drawn independently according to (6). The resulting distribution is also well fitted by an exp-gamma distribution, at least in the range where there are empirical data. This result suggests that the gamma distribution identified as a macroecological pattern [10] might just be the result of a statistical aggregation [43]. **(f)** Abundance time series retain certain turnover in this case, which might explain the different deviations between the distributions **(c)** and **(e)**. Simulations of the metacommunity (4) are run with *M* = 30, *µ* = 10*/S*, 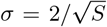, *x*_ext_ = 10^−20^, *D* = 10^−3^ and *ρ* = 0.95. Additional parameter combinations are explored in Figs. S11–S16, while alternative landscape geometries and choices of the diffusion coefficients are investigated in Figs. S17–S20. Simulations of the open model with constant migration, (2), use the same parameters, with *λ* = 10^−4^ instead of *D*. A detailed macroecological characterization of the single-patch dynamics is presented in Fig. S21.

These findings are robust across different parameter combinations, including diffusion rate, number of patches, interaction mean and variance, and inter-patch interaction correlation (see Figs. S11–S16). Robustness also extends to alternative spatial geometries of the patch distribution (Figs. S17–S19) and different choices of the diffusion coefficients (Fig. S20).

It has been recently shown [43] that gamma distributions can arise by aggregating positive random variables, as a sort of precursor of the central-limit normal behavior. It is then tempting to see the last result as an example of this effect. In order to test for this, we generated *M* realizations of the interaction parameters as in (6), and used each of them to run a simulation of the gLV model with migration, (2), starting from a random initial condition. The migration rate was set to a value similar to the diffusion constant. Then we have aggregated the abundances of all *M* runs and obtained the corresponding AFD. The results are shown in Figs. 4**e** and **f**. An exp-gamma distribution also makes a good fit to these data, as long as *P* (*z*) *>* 10^−4^, suggesting that Grilli*’*s first law (abundances are gamma-distributed) might not be revealing any mechanism of microbial communities, but rather be an effect of aggregation due to the large spatial resolution of the empirical samples.

But, as mentioned above, the aggregated distribution of the metacommunity exhibits deviations in the left tail with respect to the exp-gamma—corresponding to the lowest abundances for each species. Are they real? Interestingly, comparing the empirical data with this synthetic distribution reveals that it reproduces them more accurately than a fit to the exp-gamma distribution (root mean square error, RMSE, of the synthetic AFD: 0.4143; RMSE of the fit to an exp-gamma distribution: 2.934). The reason for this is that there are outliers at the extreme left tail of the empirical distribution that are well captured by the deviations of the aggregated distribution. Unfortunately, there are too few outliers to get any sound conclusion out of this (see, however, Figs. S9 and S10). Nevertheless, this result suggests that understanding the mechanism of microbial dynamics may require to gain an order of magnitude in the resolution of abundance data.

## V. DISCUSSION

This work has been motivated by the research question: Are fluctuations in microbial communities predominantly driven by interactions? Many relevant contributions have been made over the last decade in an attempt to answer this question. Most of them use the framework of the generalized Lotka-Volterra model, as it is the simplest possible description of an interacting ecological community. It was soon recognized that maintaining chaotic fluctuations in such a system is possible only provided it remains open to a reservoir that constantly supplies and restores extinct species. Both simulations and mean-field theory reveal that these fluctuating dynamics consist of a species turnover that constantly brings species into and out of the extant community—whose composition varies over time.

However, all these contributions share a serious drawback that prevents them from providing a convincing affirmative answer to the above research question—their predictions do not match well-known, empirical macroecological patterns. More precisely, the distribution of abundance fluctuations diverges significantly from the gamma distribution that is observed in virtually every microbial biome. It is precisely the species turnover that renders the distributions so different from empirical ones—real abundances fluctuate more steadily around their mean values over time.

Our proposal to resolve this discrepancy between theory and observations is to incorporate space into the model. On the one hand, introducing space—in the form of a metacommunity—eliminates the need to add a reservoir that replenishes *ad hoc* extinct species. Migration from patch to patch within the metacommunity largely reduces the probability that a species goes extinct in the entire metacommunity—although it does not eliminate it completely. On the other hand, every patch can maintain its own transient species composition, so that, even though each of these local communities can sustain only a small number of species, the spatially extended community exhibits a high diversity as a whole. It is not surprising that space has been proposed [49, 51] as a possible way out of the diversity-stability dilemma that May*’*s work introduced into the ecological debate [52].

In this work, we have shown that by treating the metacommunity as a whole, the fluctuations of global abundances do follow the gamma distribution observed in empirical data, even though such a pattern cannot be found in any single patch. This finding casts this macroecological pattern in a new light, showing that it might simply be the result of statistical aggregation [43] rather than the consequence of a hidden mechanism within microbial communities. Furthermore, the synthetic distribution obtained in simulations shows deviations from the gamma distribution at very low abundances, close to the resolution limit of empirical data. Remarkably, real data exhibit outliers precisely in the region where these deviations appear. Although this falls short of empirical confirmation, it is highly suggestive. All these findings are robust to changes in both the interaction parameters and the number of patches in the metacommunity.

But why should we examine aggregated, rather than local, abundances when making comparisons with empirical data? By the very nature of the sampling process, aggregation is unavoidable. Empirical samples contain massive amounts of bacteria that necessarily represent a coarse-graining of the microbial community. For instance, 1 g of soil sample can contain around 10^8^–10^10^ bacteria [53]; aquatic environments are analyzed by filtering on the order of 20–100 l of water [54], thus collapsing communities that are meters away from each other; a typical fecal sample (∼ 100–200 mg) contains around 10^10^–10^11^ bacteria [55]. If, as the metacommunity model shows, the local dynamics are very different from the global ones, they remain hidden within these coarse data.

Incidentally, the shape of the AFDs has been reported to be invariant under different taxa definitions across the phylogenetic tree [56]. This is hardly surprising if what we are observing are aggregated abundances. As a matter of fact, this seems a more plausible explanation for phylogenetic invariance than associating functional similarity to genetic relatedness—a dubious association at best if one takes into account the complexities of genotype-to-phenotype maps [57]. In the light of the present work, phylogenetic invariance appears as a further piece of evidence for the presence of aggregation in empirical data.

A reasonable question at this point is to ask about Taylor*’*s law and the lognormal distribution of mean abundances. Neither of these two patterns is captured by the results of Eq. (4) as such because the rescaling of abundances by the carrying capacities changes their mean values. However, they are automatically restored by explicitly choosing carrying capacities according to a lognormal distribution—just as in Ref. [10] (see SM, Fig. S22). Although reassuring, this result is not particularly revealing about the origin of these two patterns because it amounts to introducing a large number of parameters *ad hoc*. The ultimate cause of these patterns may lie in a different choice of the distribution of interactions [39] or as a consequence of a more sophisticated consumer-resource dynamics [58]. In any case, for the purpose of the present work, the scale chosen to represent abundances does not affect the representation of the AFD in terms of the variable *z* introduced in (5). Since studying this pattern was our main goal, scaling abundances by the carrying capacity circumvents the need to introduce many unknown parameters.

For the purpose of future research, our results suggest two lines of experimental research. One involves increasing the resolution of abundance data with the aim of capturing possible deviations from a gamma distribution. The second requires creating large-scale, well-mixed communities where samples would be representative of local abundances. This might be achieved, e.g. by stirring the reservoir containing the microbial community while ensuring at the same time that no species goes extinct. This way any species could interact with virtually any other—thus eliminating space from the equation. If the Lotka-Volterra model is a reliable description of the community dynamics, the resulting log-abundance distribution should exhibit the two-peaked structure described above.

Is spatial coarse-graining the only possible explanation for the observation of a gamma distribution of abundance fluctuations? Not necessarily. A different (open) model has very recently been proposed where species interactions emerge via resource consumption and cross-feeding [58]. This model is capable of reproducing known macroecological patterns—the gamma distribution among them—without the need to introduce spatial coarse-graining. Which model provides a more suitable description of microbial communities—or whether there are different scales at which each one operates—is a question for future experimental work.

The type of analysis developed here, together with the experiments it suggests, has implications that extend beyond the specific problem of the distribution of abundance fluctuations and the role of space, touching on several central issues in contemporary microbial ecology. First of all, it has important implications on the nature of interspecific interactions, a topic that has attracted considerable attention in recent years, particularly regarding the relative prevalence of mutualism versus competition [59, 60]. Were the dynamics here described empirically confirmed, they would indicate that interactions in microbial communities are predominantly competitive and not necessarily weak. Secondly, the scenario here described challenges the prevailing view according to which microbial ecosystems operate near a stationary state, possibly perturbed by external stochastic fluctuations. Instead, it suggests that microbial populations follow intrinsically out-of-equilibrium, chaotic trajectories—in agreement with recent theoretical insights [28]. Taken together, these perspectives call for a rethinking of both the structure of interactions and the dynamical regime of microbial communities, in order to achieve a deeper understanding of their behavior.

## VI. ACKNOWLEDGMENTS

JM Camacho, M Castro, MA Muñoz, and E Nieto read the manuscript and made helpful suggestions. MICIN/AEI/10.13039/501100011033 and *“*ERDF/EU A way of making Europe*”* funded this work under grant PID2022-141802NB-I00 (BASIC). AG-A is supported by an FPU predoctoral contract (FPU24/02195) from the Spanish Ministry of Science, Innovation and Universities (MCIU).

## Appendix A Abundance fluctuation distribution

Analysis of nine real biomes from *MGnify* (formerly EBI Metagenomics) [50] reveals that fluctuations in the abundances of different species, obtained by sampling time series or different samples of microbial communities, can all be fitted by the same distribution. Although initially identified as a gamma distribution [10], recent work shows a generalized gamma distribution fits more accurately the empirical data [61]. This distribution is parametrized as

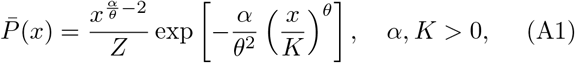

where *Z* is a normalization constant. The gamma distribution—which is the stationary state of Grilli*’*s SLM [10]—corresponds to *θ* = 1, however, other choices of this parameter account for a wider—and arguably more realistic—range of underlying stochastic dynamics, and in fact data are consistent with values of *θ* in the range 0 ≲ *θ* ≲ 5 [61].

Whichever the value of *θ*, abundances from different samples—which vary across several orders of magnitude—can only be compared by using the standard variable *z* defined in (5). If *x* follows the distribution in (A1), *z* follows the exp-gamma distribution

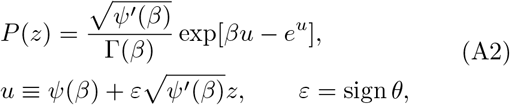

where *β* = (*α* − *θ*)*/θ*^2^ and *ψ*(*x*) = Γ^′^(*x*)*/*Γ(*x*) denotes the digamma function. As this distribution depends only on the parameter *β* and the sign of *θ*, it cannot distinguish between a gamma and a generalized gamma with *θ >* 0. If *θ <* 0 though, it has the opposite skewness. Figure S1 of the Supplementary Information (SI) shows that empirical data taken from very diverse biomes can all be fitted by the distribution in (A2) with *ε* = +1.

## Appendix B Numerical integration of the equations

Simulations of the open Lotka-Volterra model with constant migration, (2), were integrated numerically using the ode15s solver implemented in MATLAB, based on variable-order backward differentiation formulas (BDFs) [62], which is particularly suitable for stiff systems, given that strong interactions, large community sizes, and small migration rates can generate widely separated time scales. Simulations were carried out on the variables *z*_*i*_(*t*) = log *x*_*i*_(*t*) in order to guarantee positivity of the abundances. Simulations of the metacommunity model, (4), were instead performed in Python using the solve ivp routine with a Runge–Kutta solver, applied directly to the abundances *x*^*u*^(*t*); in this case, the log-transformation is unnecessary since the extinction threshold *x*_ext_ prevents abundances from reaching zero. In both cases, running times go up to *t* = 30000 with a time step Δ*t* = 0.1. To ensure that SADs and AFDs are calculated in the stationary state, we discarded all data for *t* ≤ 500 and performed calculations with the remainder of the time series. Unless otherwise specified, simulations were initialized with homogeneous abundances *x*_*i*_(0) ∼ *U*= 0.5 for all species. We verified that sampling initial conditions from a uniform distribution *x*_*i*_(0) [0, 1] leads to the same qualitative dynamical behavior and statistical patterns, indicating that the results are robust with respect to the choice of initial conditions.

## Appendix C Curve fitting

Fits of the analytical distributions (exp-gamma and Student*’*s *t*) to empirical or synthetic AFDs are performed on log *P* (*z*) rather than on *P* (*z*) itself, in order to properly capture the behavior of the tails. When comparing with empirical data, as in Fig. 4 and Figs. S11– S16, the fitting procedure is restricted to the interval −5 ≤ *z* ≤ 5, corresponding to the range of fluctuations sampled in the empirical biomes.

## Appendix D Trajectory classification

Simulated trajectories can be classified as *‘*failed*’, ‘*stable*’* or *‘*fluctuating*’*, and the latter in turn as *‘*periodic*’* or *‘*aperiodic*’*. Trajectories for which the numerical integration does not complete, corresponding to unbounded growth dynamics, are classified as failed. For the remaining trajectories, we first calculate the coefficient of variation (standard deviation/mean) of the time series after discarding the initial transient. If the average coefficient is below (above) 0.01, the trajectory is considered stable (fluctuating). We then compute the fast Fourier transform (FFT) of the fluctuating trajectories, 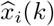, and normalize it by its first positive-frequency component, 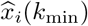, where *k*_min_ *>* 0 denotes the smallest discrete frequency, in order to make Fourier modes comparable across species and enhance signal detection. Finally, as all abundances share the same periodicities, we compute

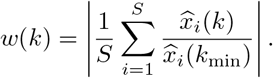

In communities with periodic dynamics, *w*(*k*) exhibits sharp spikes (typical height ∼ 10^2^) at the fundamental frequency and its harmonics; in communities with aperiodic dynamics, by contrast, *w*(*k*) remains much smaller and does not display such pronounced peaks. This allows us to distinguish periodic from aperiodic dynamics in an automated manner. In practice, we classify fluctuating trajectories as periodic when max_*k>*0_ *w*(*k*) *>* 10, and as aperiodic otherwise. Figures S23 and S24 of the SI show examples of periodic and aperiodic time series, together with the corresponding *w*(*k*).

