## Supplementary Material for "The role of space in explaining macroecological patterns of microbial abundance"

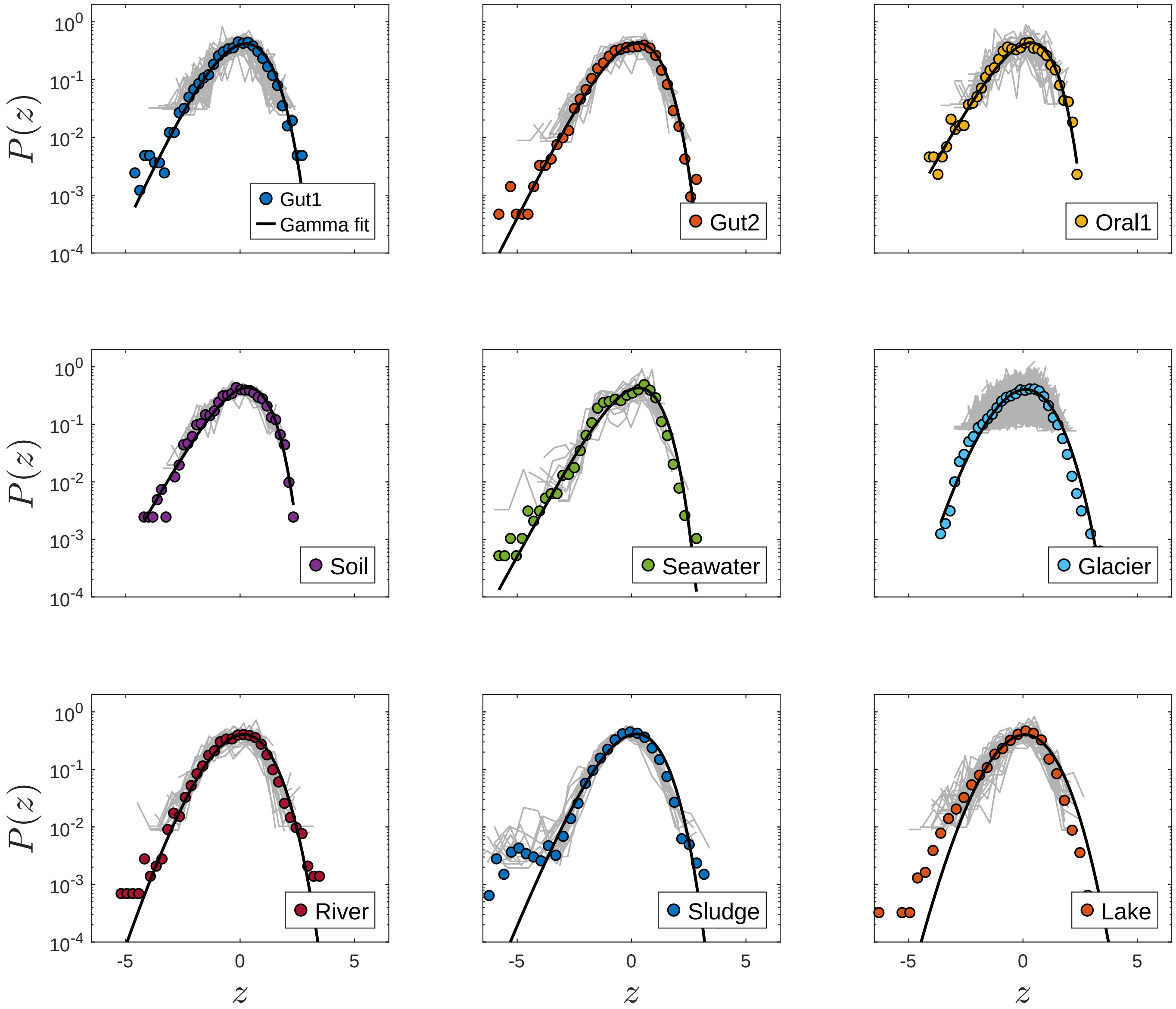


FIG. S1. **Empirical abundance fluctuation distributions.** Distribution of abundance fluctuations for nine real biomes obtained from the *MGnify* platform [1]. For each biome, we select species present in all samples and consider the logarithm of their relative abundances, defined as the ratio between read counts and the total number of reads. These values are then rescaled according to the standardized variable $z$ defined in Eq. (5) of the main text, and the resulting species-wise distributions are shown as solid grey lines. Coloured dotted lines represent the distributions obtained by aggregating relative abundances across all species within each biome. These aggregated distributions are well fitted by the exp-gamma distribution [Eq. (8) of the main text] with $\varepsilon = +1$ (black solid line). This distribution corresponds to a gamma distribution expressed in terms of $z$.

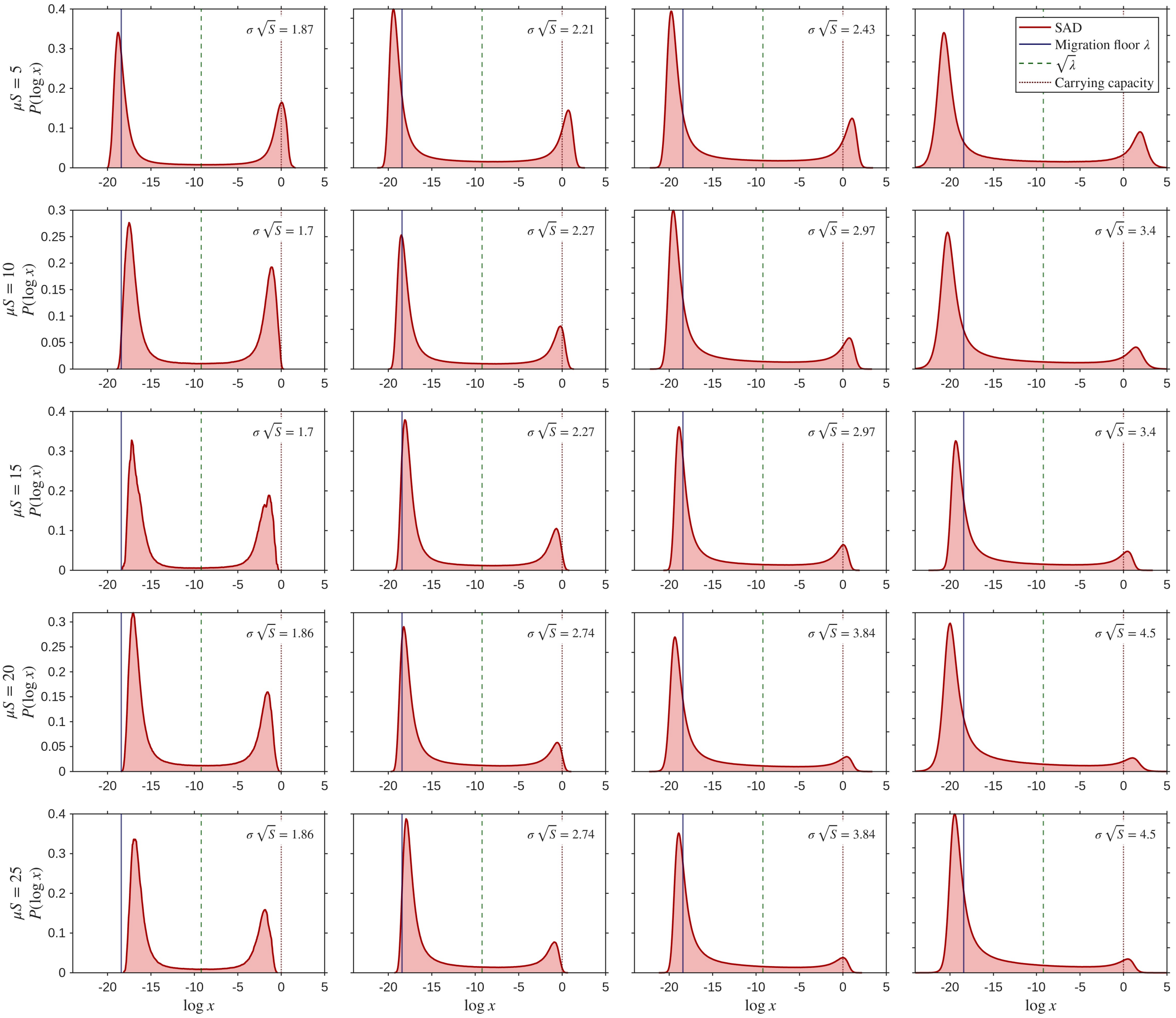


FIG. S2. **Species abundance distribution as a function of interaction parameters.** Distribution of species abundances for the open Lotka-Volterra model with constant migration [Eq. (2) of the main text]. Panels correspond to different values of the interaction mean $\mu$ (rows) and standard deviation $\sigma$ (columns), within the multiple-attractor phase where chaotic turnover is observed. The resulting distributions are bimodal, with peak heights and positions varying across parameter values, while remaining centered around the migration floor and carrying capacity, indicated by the vertical dashed lines. Simulations were performed with $\lambda = 10^{-8}$, $S = 1000$. Abundances were sampled every $\Delta t = 0.1$ after a transient, over the time window $500 \leq t \leq 15000$.

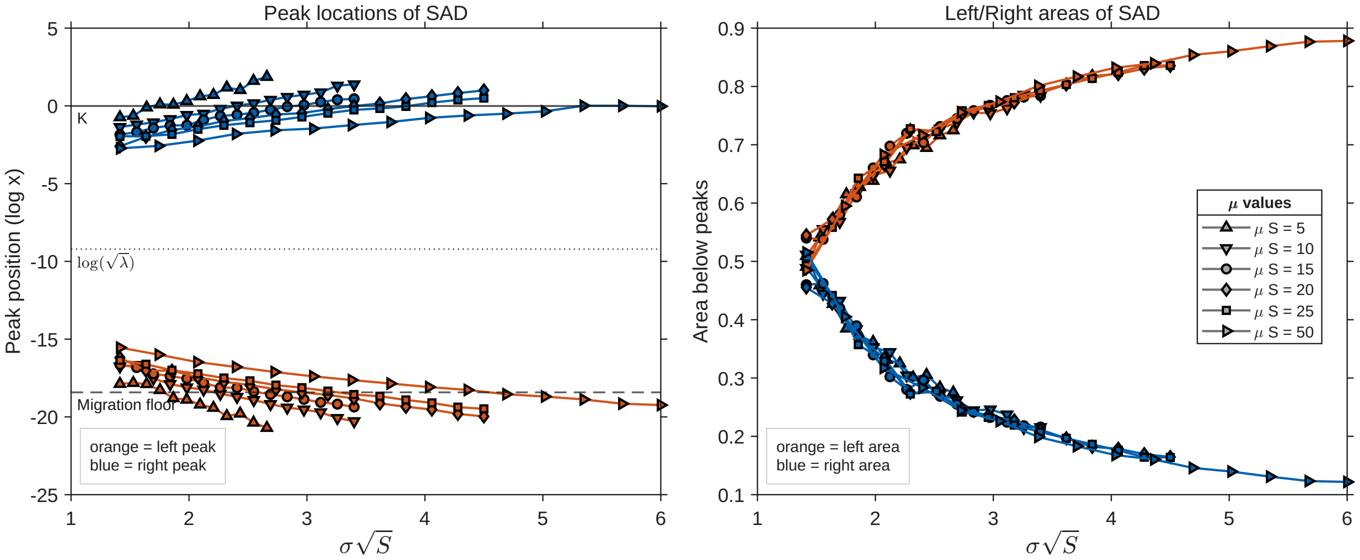


FIG. S3. **Peak positions and areas of the species abundance distribution as a function of the interaction parameters.** **(a)** Position of the left (orange) and right peaks (defined as in Fig. 3**d** of the main text) as a function of the standard deviation of the interactions $\sigma$, for different values of the mean interaction strength $\mu$, within the multiple-attractor phase ($\sigma\sqrt{S} > \sqrt{2}$). As $\mu$ increases, the two peaks move slightly closer to each other, whereas increasing $\sigma$ shifts them farther apart, as can also be appreciated in Fig. S2. For reference, the carrying capacity and the migration floor, which set the characteristic centers of the two peaks, are also indicated. **(b)** Peak areas as a function of $\sigma$, showing how the relative weight of the two modes changes across parameter values. They depend weakly on $\mu$. The simulation parameters are the same as in Fig. S2.

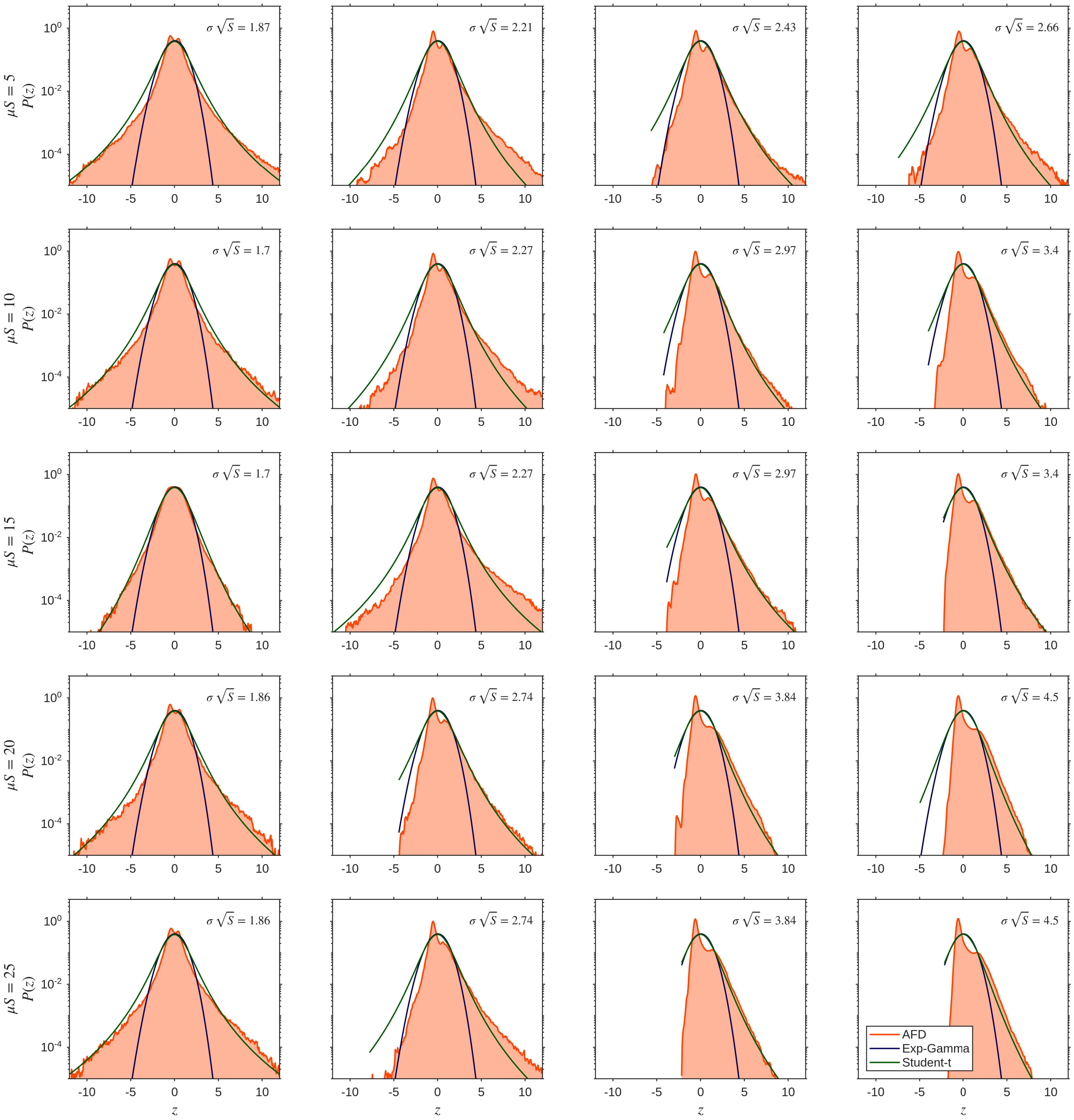


FIG. S4. **Abundance fluctuation distribution as a function of interaction parameters.** Distribution of fluctuations of the standardized log-abundance $z$ defined in Eq. (5) of the main text, for the open Lotka-Volterra model with constant migration [Eq. (2)], aggregated across all species in the community. Panels correspond to different values of the standard deviation of the interactions $\sigma$ and the mean interaction strength $\mu$, within the multiple attractors phase. The resulting distributions deviate markedly from a gamma, with tails better described by a Student's $t$ distribution and a bimodal structure associated with turnover dynamics. Simulations were performed with $\lambda = 10^{-8}$, $S = 1000$. Abundances were sampled every $\Delta t = 0.1$ after discarding the transient and retaining the interval $500 \leq t \leq 15000$.

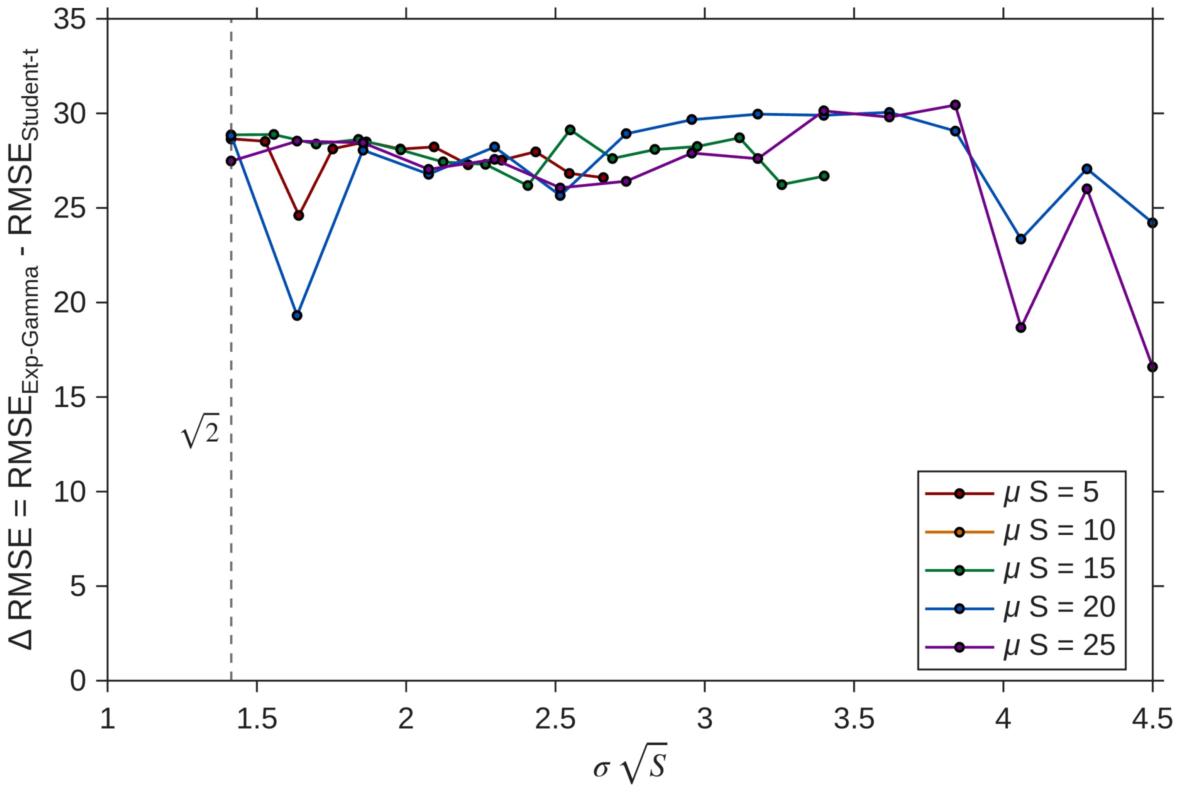


FIG. S5. **Distance of the abundance fluctuation distribution (AFD) from fitted functions.** Root-mean-square error (RMSE) between the AFD obtained for the open Lotka-Volterra model with constant migration [Eq. (2) of the main text], with representative examples shown in Fig. S2, and two candidate fitting curves: an exp-gamma distribution and a Student's $t$ distribution. The RMSE is plotted as a function of the standard deviation of the interactions $\sigma$, for different values of the mean interaction strength $\mu$. Across the explored parameter range, the RMSE of the exp-gamma fit is systematically one order of magnitude larger than that of the Student's $t$ fit. The vertical dashed line refers to $\sigma_c\sqrt{S} = \sqrt{2}$, corresponding to the phase-transition threshold into the multiple-attractor phase of the generalized well-mixed Lotka-Volterra model, as shown in [2]. Both the RMSE and the least-squares fitting procedure were computed using $\log P(z)$ and the logarithm of the fitted distributions, in order to better capture deviations in the tails. All simulation parameters are the same as in Fig. S4.

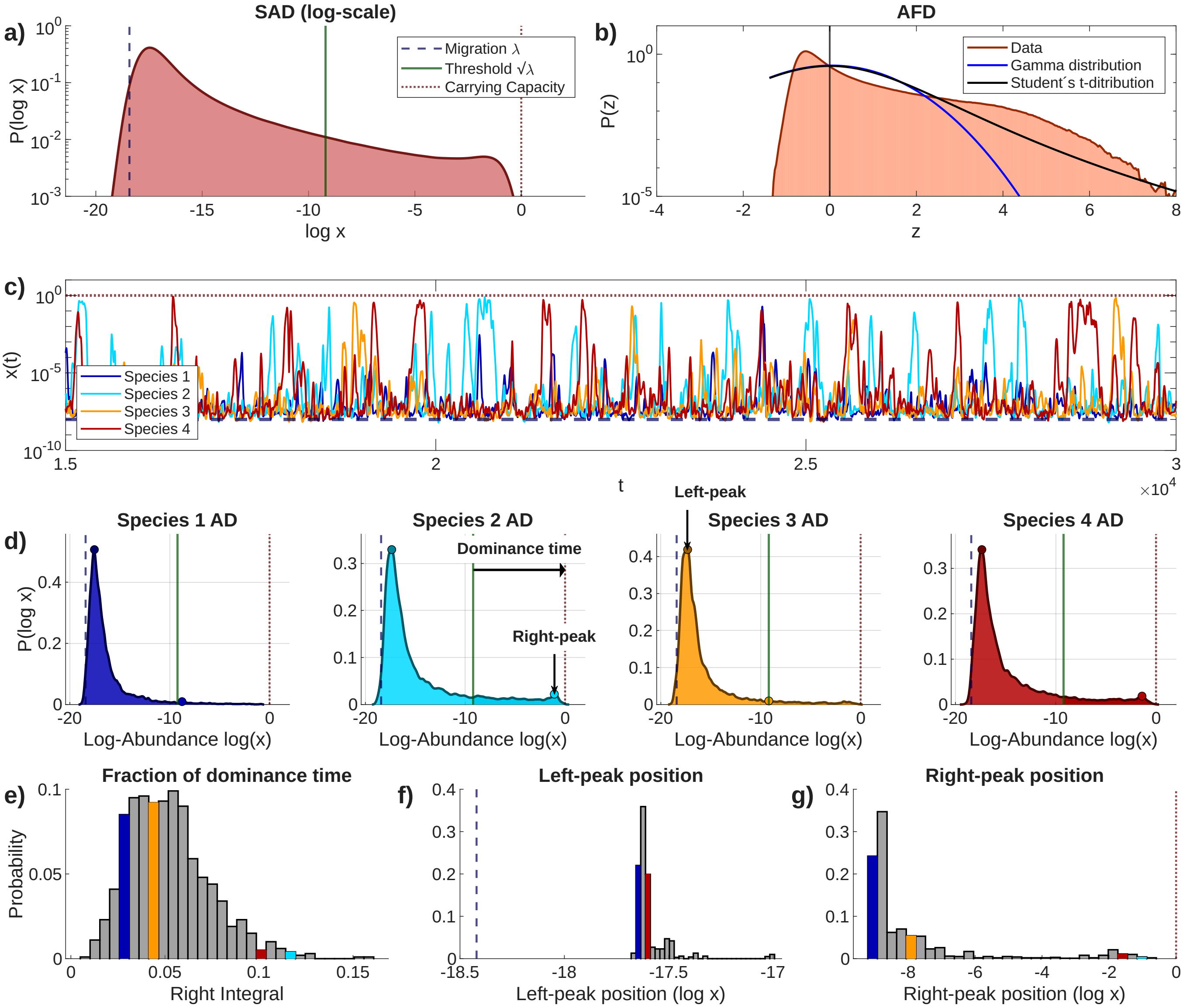


FIG. S6. **Macroecological characterization of the open Lotka-Volterra with migration in the strong-interaction regime.** **(a)** Species log-abundance distribution (distribution of log-abundances across time samples), aggregated over all species in the community. The distribution exhibits a characteristic bimodal structure with an intermediate plateau. In contrast to Fig. 3 of the main text, the vertical axis is shown on a log scale in order to highlight the right peak, since for this parameter set the fraction of dominant species is only $10/500 = 0.02$. **(b)** Abundance fluctuation distribution (AFD), defined as the distribution of standardized log-abundances [Eq. (5) of the main text], aggregated across all species in the community. Solid curves show fits to both the exp-gamma distribution and the Student's $t$ distribution. Although the Student's $t$ distribution provides a better description than the exp-gamma, neither fit adequately captures the AFD. **(c)** Representative abundance time series for selected species. Abundances fluctuate between the migration floor $\lambda$ and the carrying capacity, displaying heterogeneous temporal behaviors ranging from small fluctuations to rapid turnover between rare and dominant states. **(d)** Log-abundance distributions for the species shown in **(c)**. **(e)** Histogram of dominance times, defined as the fraction of time during which species abundances $x_i \geq \sqrt{\lambda}$. Lower net competition is associated with a stronger dominance bias [3]. **(f), (g)** Distributions of the positions of the left and right peaks of the abundance distributions shown in **(d)**, respectively. Insets in **(d)** illustrate the definition of these quantities. Coloured bars indicate the locations of the species highlighted in **(c)** and **(d)** within the corresponding histograms. Results are obtained from simulations of Eq. (2) of the main text, with $S = 500$, $\mu = 0.5$, $\sigma = 0.3$, $\lambda = 10^{-8}$. This parameter set corresponds to the strong-interaction regime considered in [3], in which the moments of the interaction matrix are not scaled with the community size $S$, and for which a power-law species abundance distribution $P(x) \sim x^{-\nu}$ with exponent $\nu \approx 1.8$ was reported, in agreement with our simulations.

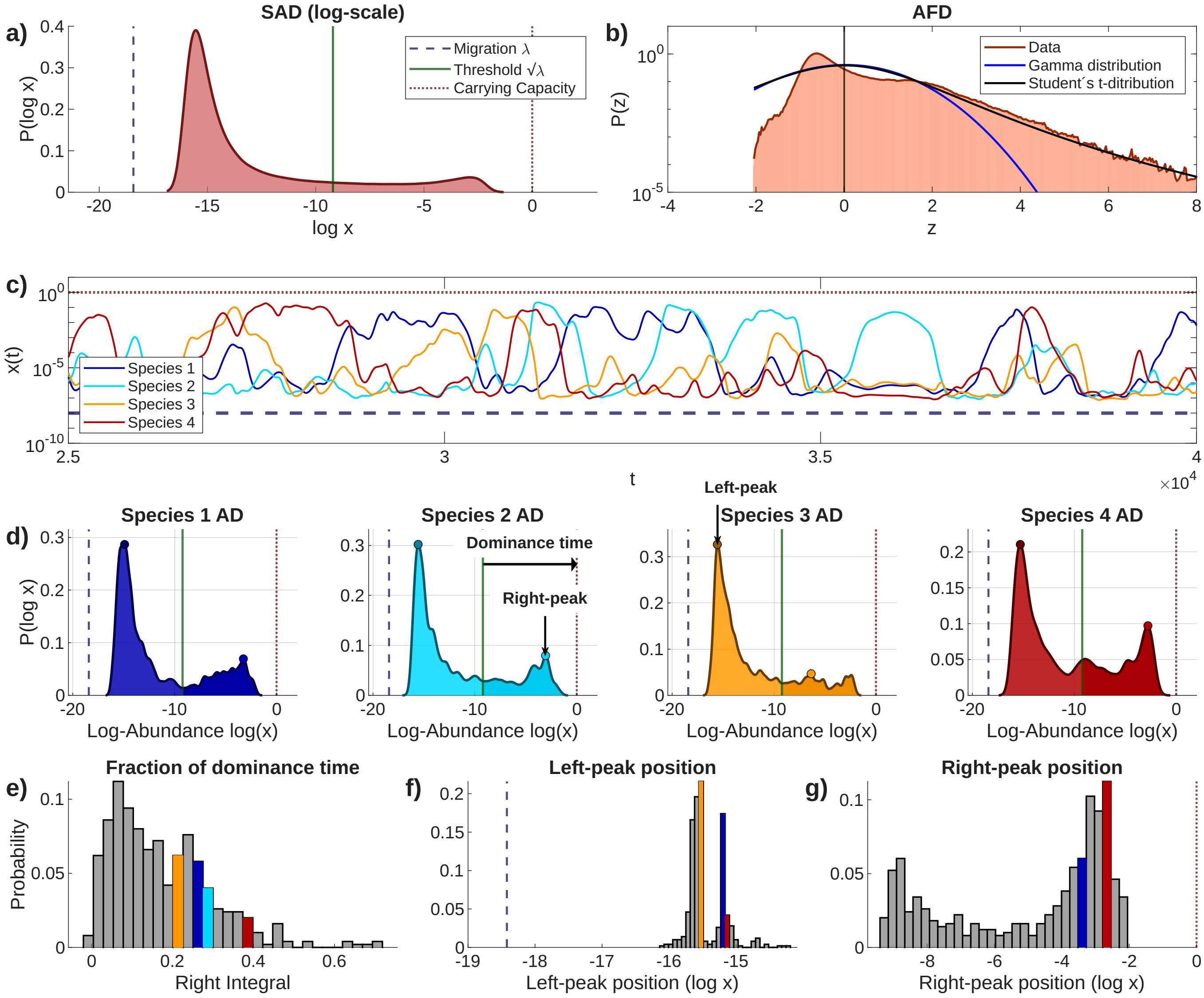


FIG. S7. **Macroecological characterization of the open Lotka-Volterra with migration in the strong-interaction regime.** Same simulations and analysis as in Fig. S6, but with a lower standard deviation of the interactions $\sigma$, so that the dominant community is more populated and the correspondence between the weak- and strong-interaction regimes becomes more apparent. **(a)** Species log-abundance distribution (distribution of log-abundances across time samples), aggregated over all species in the community. It displays a characteristic bimodal structure with an intermediate plateau. **(b)** Abundance fluctuation distribution (AFD), defined as the distribution of standardized log-abundances. Solid curves show fits to both the exp-gamma distribution and the Student's $t$ distribution. The exp-gamma fit performs markedly worse than the Student's $t$ fit. **(c)** Representative abundance time series for selected species. Abundances fluctuate between the migration floor $\lambda$ and the carrying capacity, displaying heterogeneous temporal behaviors ranging from small fluctuations to rapid turnover between rare and dominant states. Compared with Fig. S6, the lower value of $\sigma$ yields a more populated dominant community and a broader diversity of temporal behaviors. **(d)** Log-abundance distributions for the species shown in **(c)**. These distributions follow the general pattern shown in **(a)**, but the height and location of the peaks varies, reflecting a diversity of behaviors ranging from rare presence to high dominance. **(e)** Histogram of dominance times, defined as the fraction of time during which species abundances $x_i \geq \sqrt{\lambda}$. **(f), (g)** Distributions of the positions of the left and right peaks of the abundance distributions shown in **(d)**, respectively. Insets in **(d)** illustrate the definition of these quantities. Coloured bars indicate the locations in the histogram of the species illustrated in **(c)**, **(d)**. Results are obtained from simulations of Eq. (2) of the main text, with $S = 500$, $\mu = 0.5$, $\sigma = 0.1$, and $\lambda = 10^{-8}$.

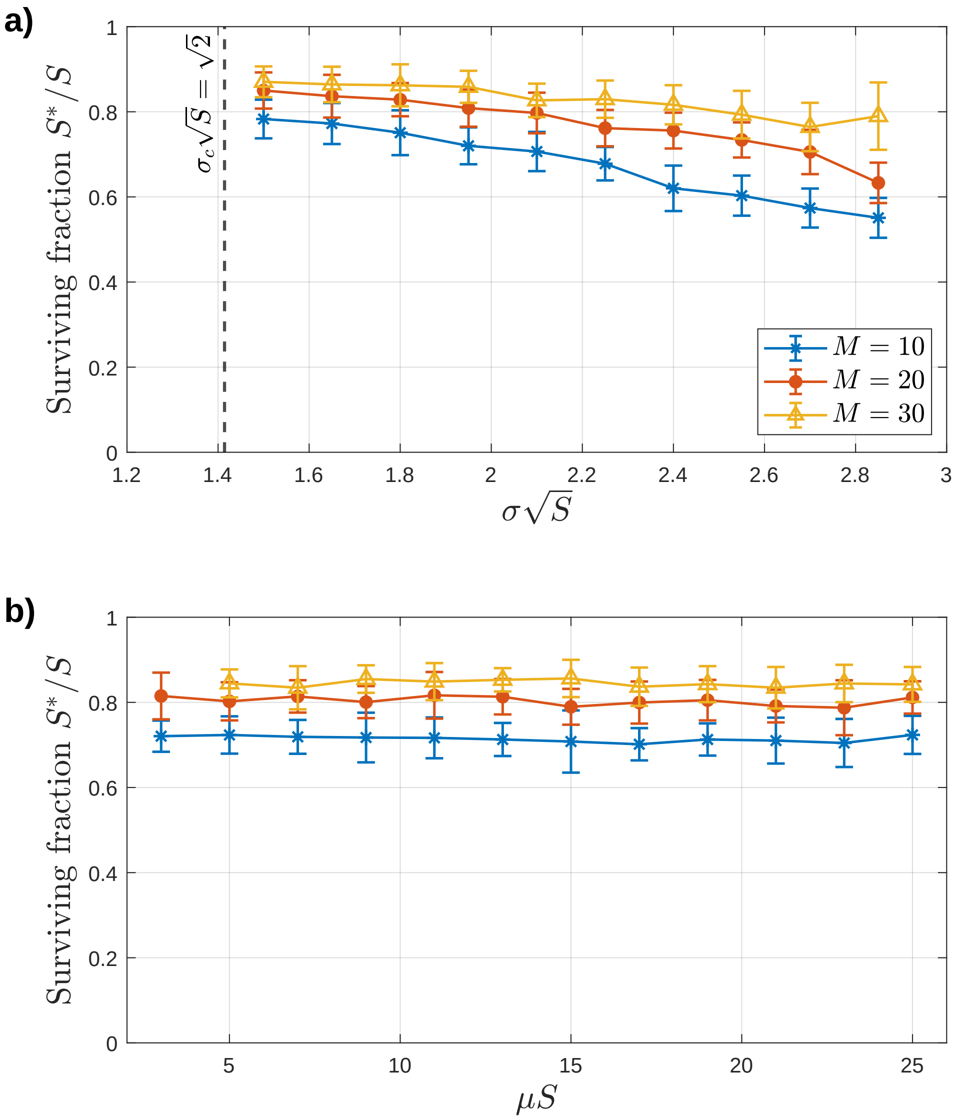


FIG. S8. **Surviving species as a function of system parameters.** Fraction of surviving species, $S^*/S$, i.e., fraction of species such that $x_i^u > x_{\rm ext}$ in all patches $u = 1, \ldots, M$, as a function of the rescaled standard deviation of **(a)** the interactions $\sigma\sqrt{S}$ and **(b)** the mean interaction strength, for different values of the number of patches $M$. Points represent the average value of $S^*/S$ over 20 realizations of the metacommunity dynamics [Eq. (3) of the main text], while error bars indicate the corresponding standard deviation. A species is considered extinct if its abundance falls below the threshold $x_{\rm ext} = 10^{-20}$ in all $M$ patches. The remaining parameters are set to $\mu S = 10$, $D = 10^{-4}$, $\rho = 0.95$, and $S = 250$. Here, $\sigma_c$ denotes the stability threshold of the generalized well-mixed Lotka-Volterra model, as shown in [2], beyond which abundances exhibit chaotic dynamics. The fraction of surviving species decreases as the interaction strength $\sigma\sqrt{S}$ or the number of patches $M$ increase, and remains nearly constant as a function of the mean interaction strength $\mu S$.

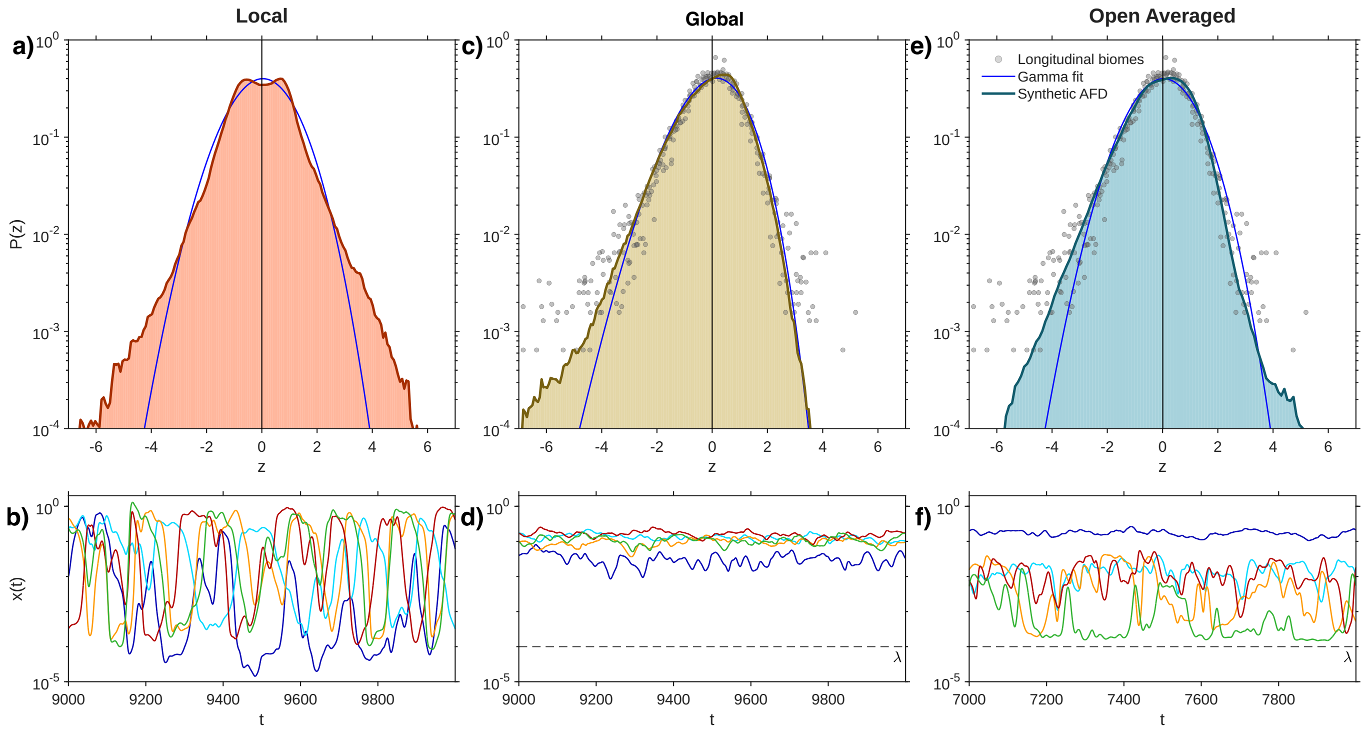


FIG. S9. **Abundance fluctuation patterns in the metacommunity model compared with longitudinal data.** Same simulations and parameters as in Fig. 4 of the main text, here compared with the empirical abundance fluctuation distributions obtained from longitudinal biome time series. **(a)** Distribution of rescaled log-abundances at the local scale, (in a single patch). The exp-gamma distribution, given by Eq. (8) of the main text, clearly provides a poor fit to the numerical distribution. **(b)** Example time series of local abundances for the same selected species as in Fig. 4**(b)** of the main text, again showing a clear turnover dynamics. **(c)** Abundance fluctuation distribution at the global scale, obtained by aggregating species abundances across all patches. Apart from small deviations in the left tail, the resulting distribution is well described by a gamma distribution. Grey dots represent empirical biome distributions from *longitudinal* data (taken from the *MGnify* platform [1]), i.e., abundances sampled over time, as in the numerical simulations. **(d)** Corresponding global abundance time series for the same selected species. In contrast to the local dynamics, these trajectories no longer exhibit turnover; instead, they show rapid fluctuations around a characteristic mean value, similar to those observed in empirical time-series. **(e)** Abundance fluctuation distribution obtained by aggregating $M$ independent realizations of the constant-migration open Lotka-Volterra model, but now with $\lambda \approx D\bar{x}_i$ consistently with the mean-field connection between the two models discussed in the *Open Lotka-Volterra models* section of the main text. Realizations are generated using different initial conditions and considering both identical and distinct interaction matrices. Blue line represents the fit to a gamma distribution. **(f)** This time, the corresponding time series still retain some turnover—although weaker and only for some species.

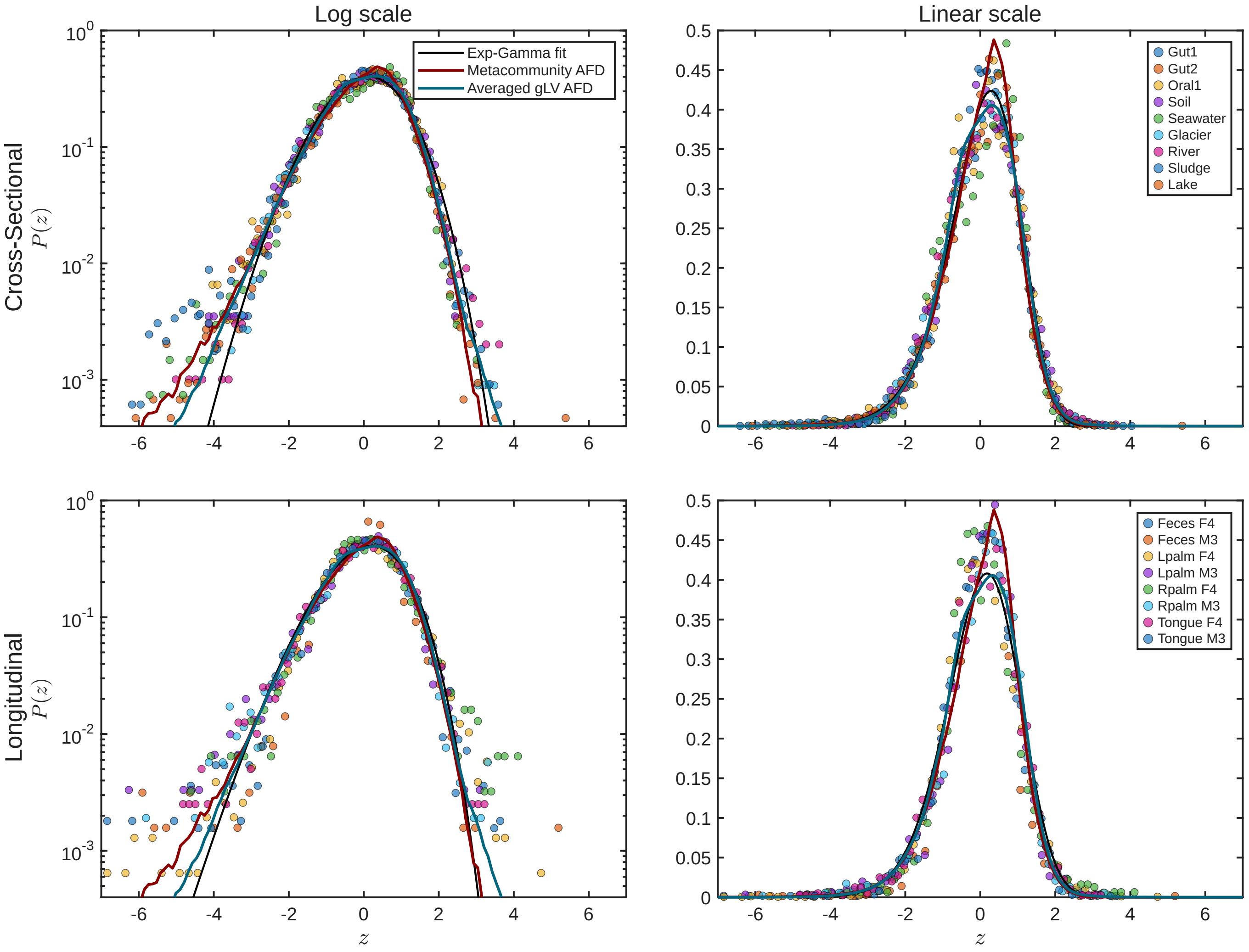


FIG. S10. **Model comparison with cross-sectional and longitudinal data.** Distribution of rescaled log-relative abundances obtained by aggregating patch dynamics in the metacommunity model (magenta solid line) and by averaging independent realizations of the gLV model with constant migration (cyan solid line), corresponding to the same cases shown in Fig. 4 of the main text. The black solid line denotes the exp-gamma distribution [Eq. (8) of the main text] fitted to the empirical AFD obtained by aggregating all biomes. Coloured dots represent empirical biome data from cross-sectional and longitudinal samples from the *MGnify* platform [1]. The probability distribution $P(z)$ is displayed on both linear and logarithmic scales. The deviation of the empirical AFD from the gamma distribution, in both longitudinal and cross-sectional samples, is well captured by the metacommunity model AFD.

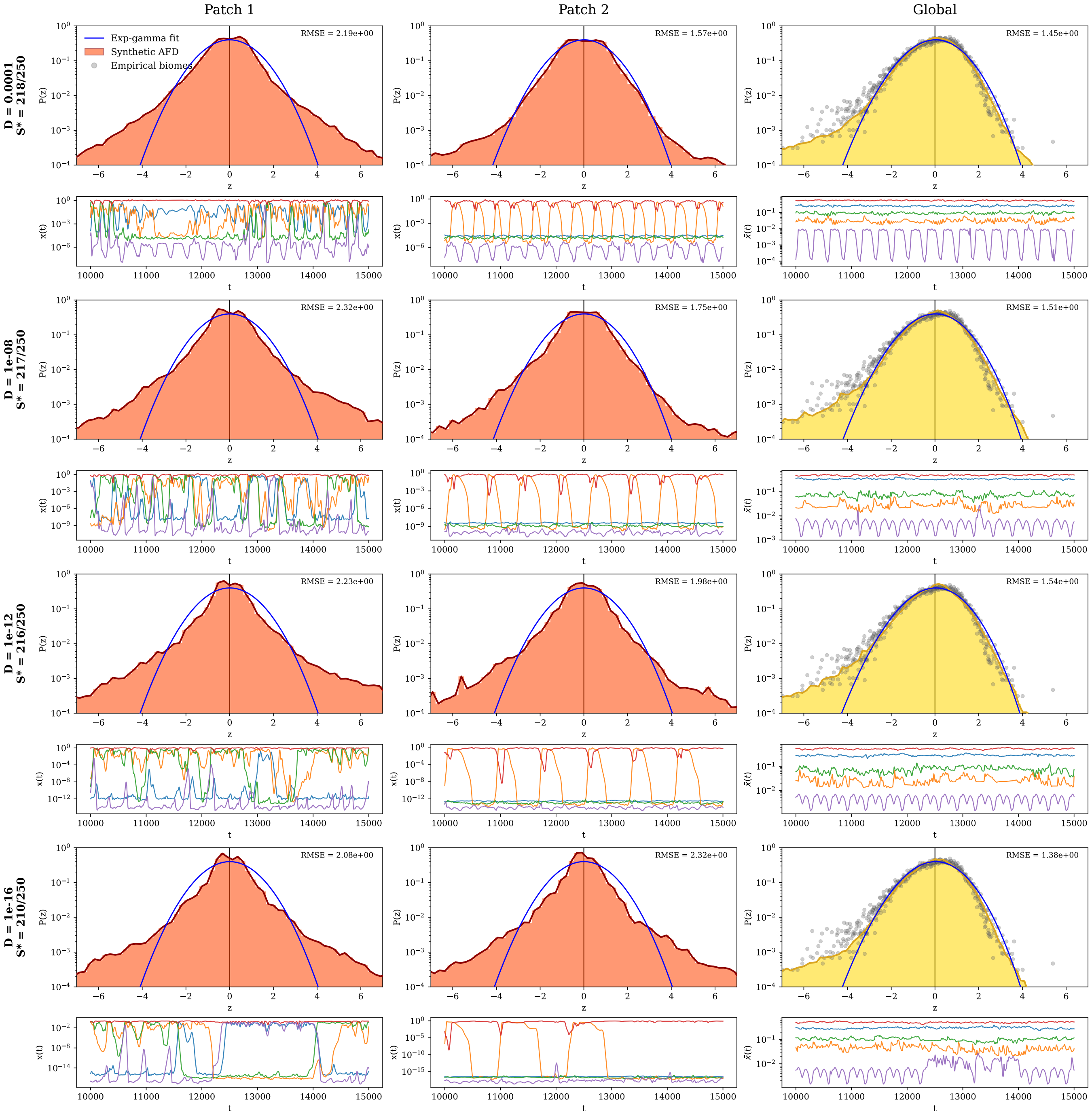


FIG. S11. **Abundance fluctuation distributions in the metacommunity model for different migration rates.** Distributions of rescaled log-abundances and the corresponding time series are shown at both the local scale (for two representative patches) and the global scale, i.e. after aggregating abundances across patches, as illustrated in Fig. 4 of the main text. As in that figure, local abundance distributions deviate significantly from the exp-gamma distribution; the root mean square errors (RMSE, reported in the top-right corner of each panel) are consistently larger for local than for global abundances. In contrast, the aggregated distributions closely match those observed in empirical biomes (gray dots). Importantly, the global distribution also captures the low-abundance deviations observed in empirical data, which are not reproduced by the exp-gamma fit. Note that varying $D$ effectively shifts the migration floor of the local patch dynamics, which, consistently with the mean-field connection discussed in the main text, is set by $\lambda \approx D\bar{x}_i$. Distributions are obtained from simulations of the metacommunity model with $S = 250$, $M = 25$, $\mu S = 10$, $\sigma\sqrt{S} = 2$, $\rho = 0.95$, and $x_{\text{ext}} = 10^{-20}$.

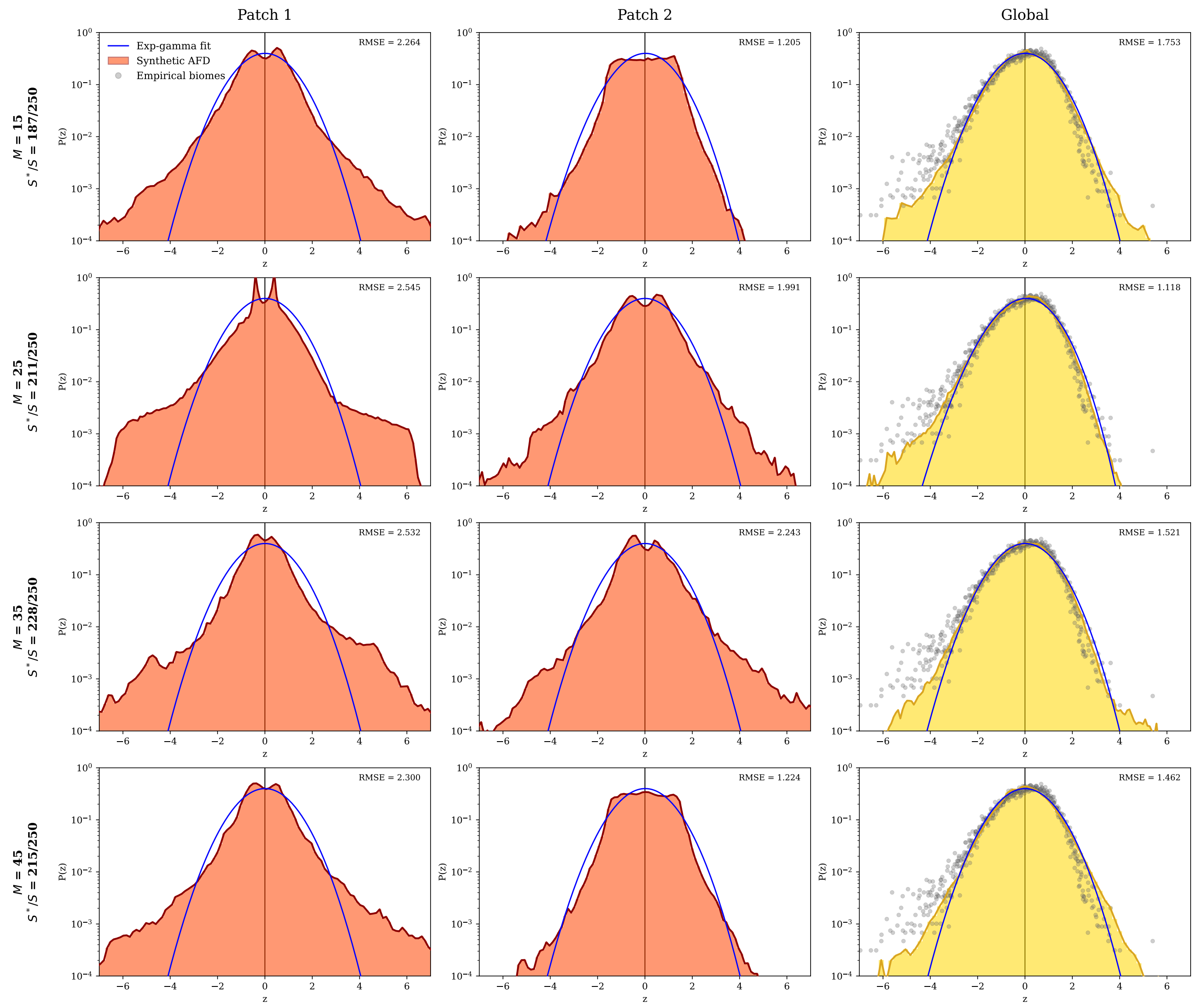


FIG. S12. **Abundance fluctuation distributions in the metacommunity model for different patch numbers.** Same as Fig. S11 but changing the number of patches $M$. Distributions are obtained from simulations of the metacommunity model with $S = 250$, $D = 10^{-6}$, $\mu S = 10$, $\sigma\sqrt{S} = 2$, $\rho = 0.95$, and $x_{\rm ext} = 10^{-20}$.

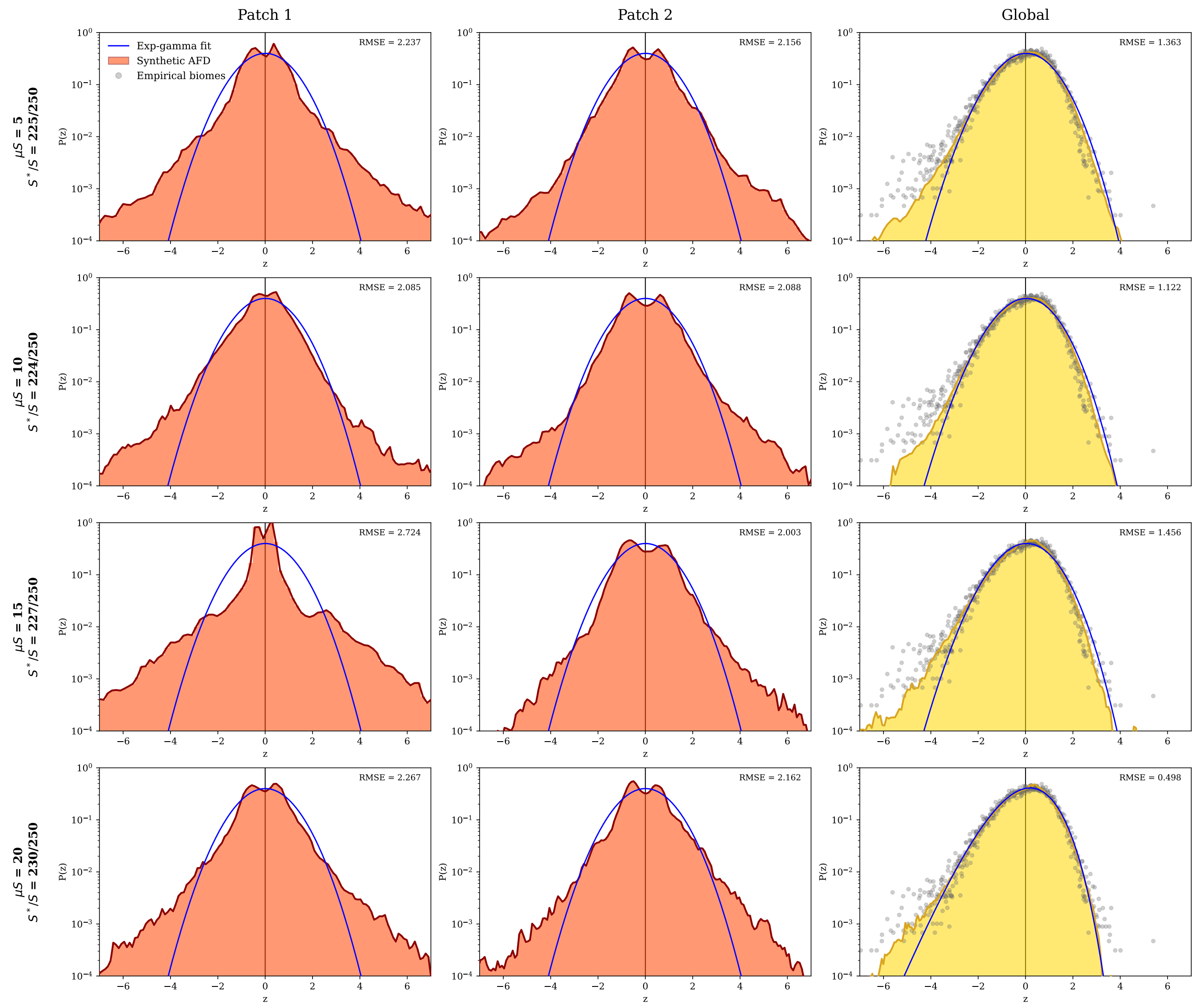


FIG. S13. **Abundance fluctuation distributions in the metacommunity model for different mean interaction values.** Same as Figs. S11 and S12 but changing the mean of the interactions $\mu S$. Distributions are obtained from simulations of the metacommunity model with $S = 250$, $D = 10^{-6}$, $M = 25$, $\sigma\sqrt{S} = 2$, $\rho = 0.95$, and $x_{\mathrm{ext}} = 10^{-20}$.

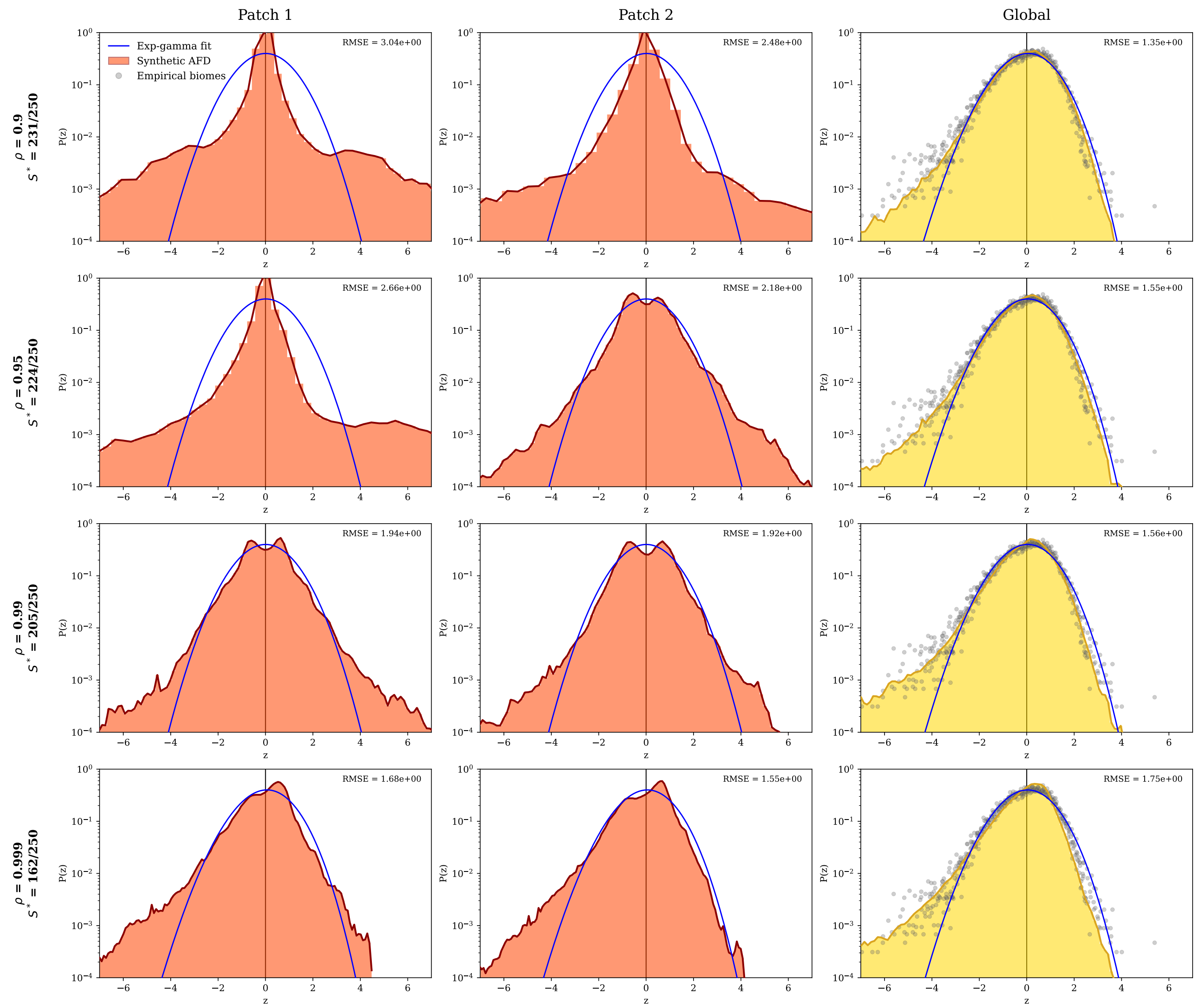


FIG. S14. **Abundance fluctuation distributions in the metacommunity model for different correlations between interaction matrices across patches.** Same as Figs. S11–S13 but varying the correlation parameter $\rho$ between interaction matrices across patches. Distributions are obtained from simulations of the metacommunity model with $S = 250$, $D = 10^{-6}$, $M = 25$, $\sigma\sqrt{S} = 2$, $\mu S = 10$, and $x_{\text{ext}} = 10^{-20}$.

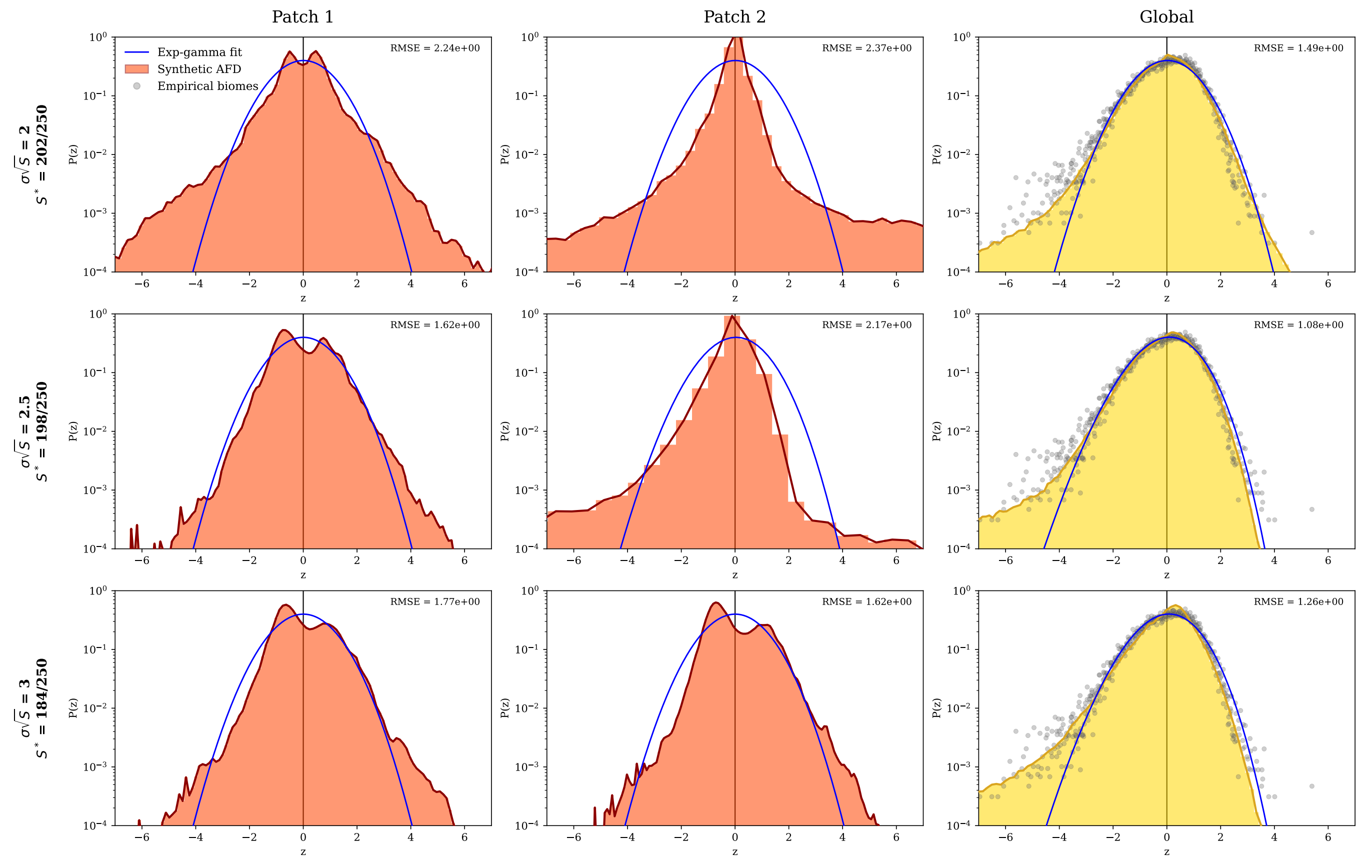


FIG. S15. **Abundance fluctuation distributions in the metacommunity model for different values of the interaction standard deviation.** Same as Figs. S11–S14 but changing the standard deviation of the interactions $\sigma\sqrt{S}$. Distributions are obtained from simulations of the metacommunity model with $S = 250$, $D = 10^{-6}$, $M = 25$, $\rho = 0.95$, $\mu S = 10$, and $x_{\rm ext} = 10^{-20}$.

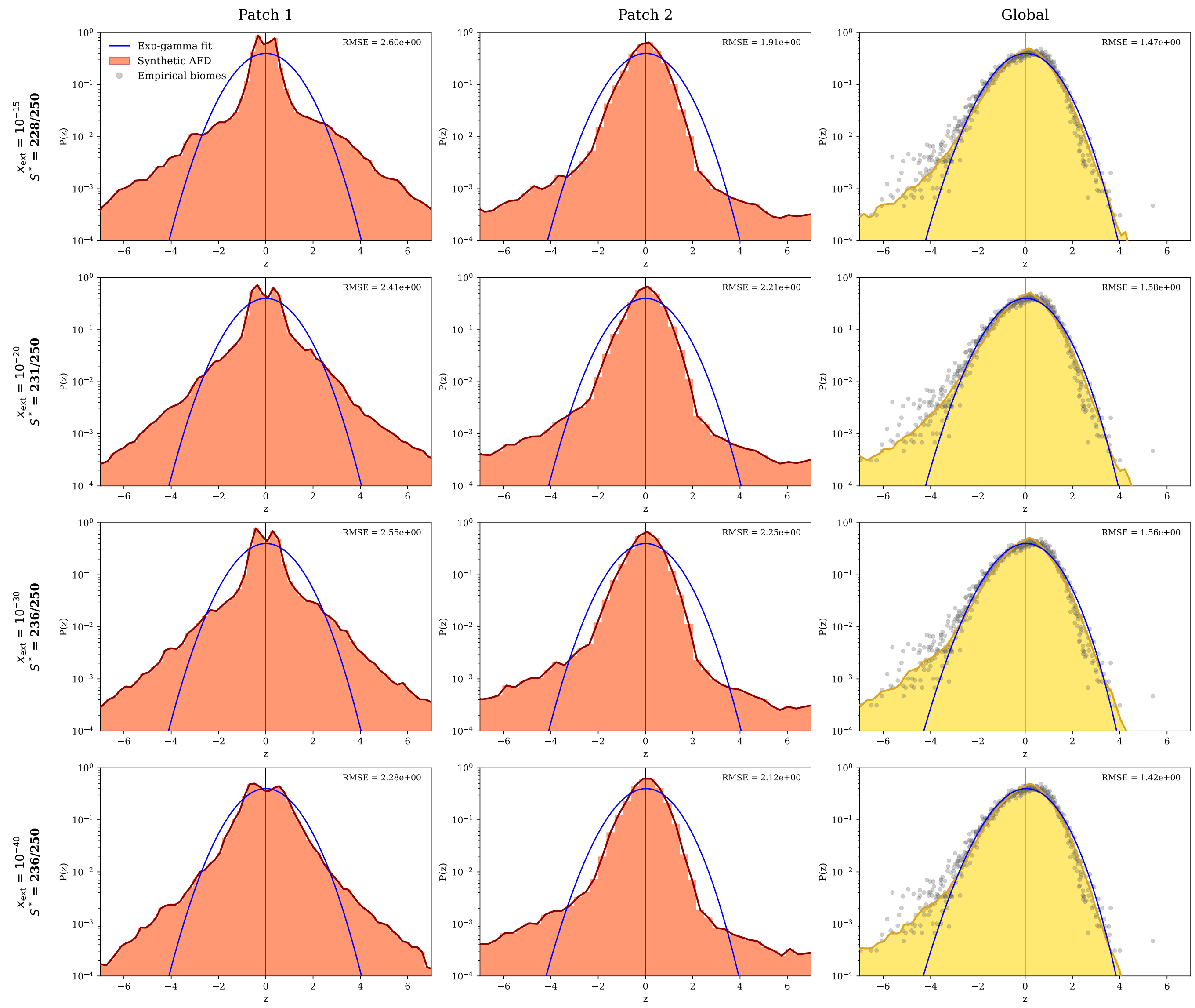


FIG. S16. **Abundance fluctuation distributions in the metacommunity model for different values of the extinction threshold.** Same as Figs. S11–S15 but changing the extinction threshold $x_{\text{ext}}$. Distributions are obtained from simulations of the metacommunity model with $S = 250$, $D = 10^{-6}$, $M = 25$, $\rho = 0.95$, $\mu S = 10$, and $\sigma\sqrt{S} = 2$.

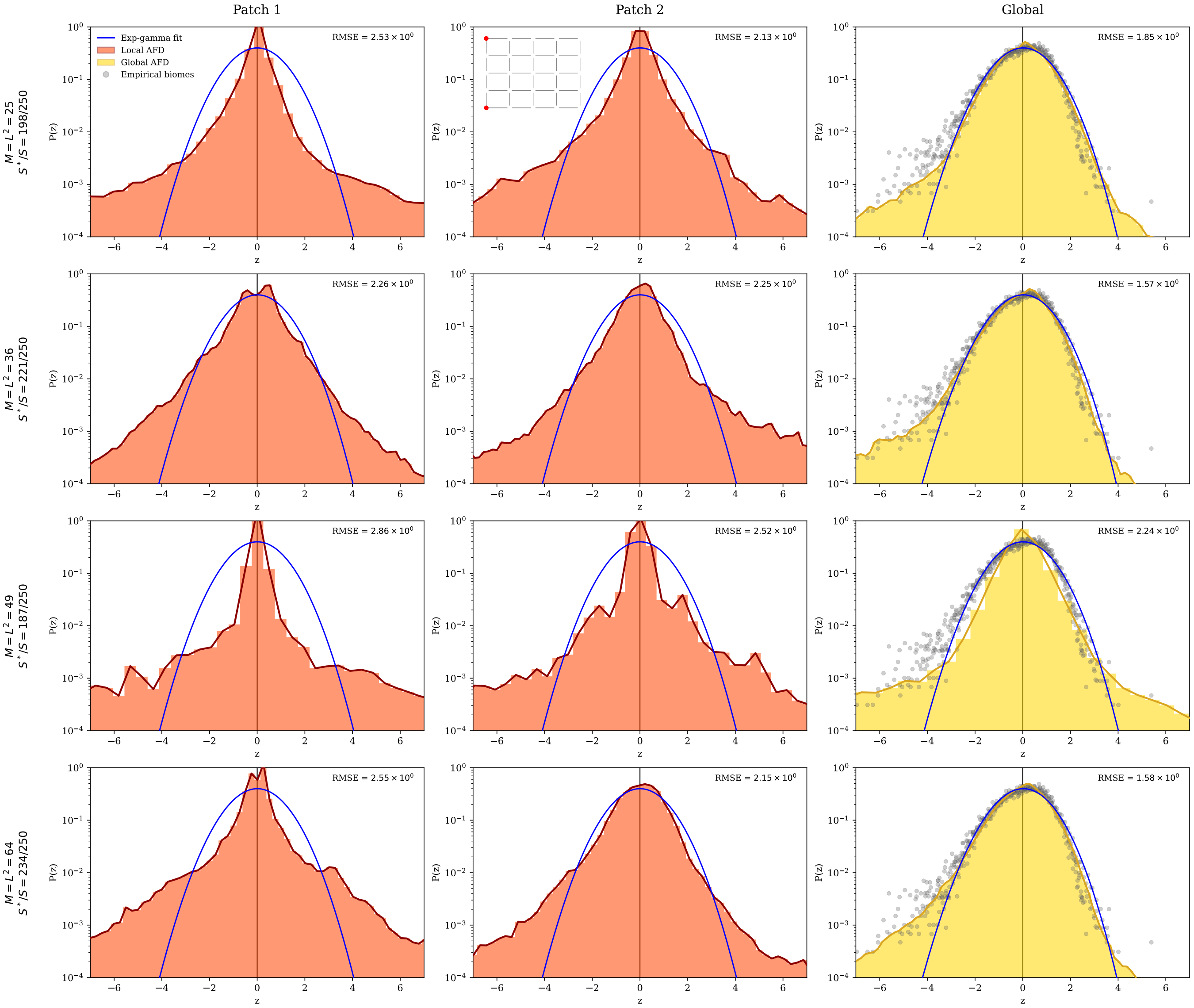


FIG. S17. **Abundance fluctuation distributions in a metacommunity with square-lattice embedding.** Same as Figs. S11–16, but with patches arranged on a square lattice. Patches are treated as vertices of the lattice, whose size $M$ varies across figure rows. Simulations are performed with $S = 250$, $D = 10^{-6}$, $\rho = 0.95$, $\mu S = 10$, $\sigma\sqrt{S} = 2$, and $\rho = 0.95$.

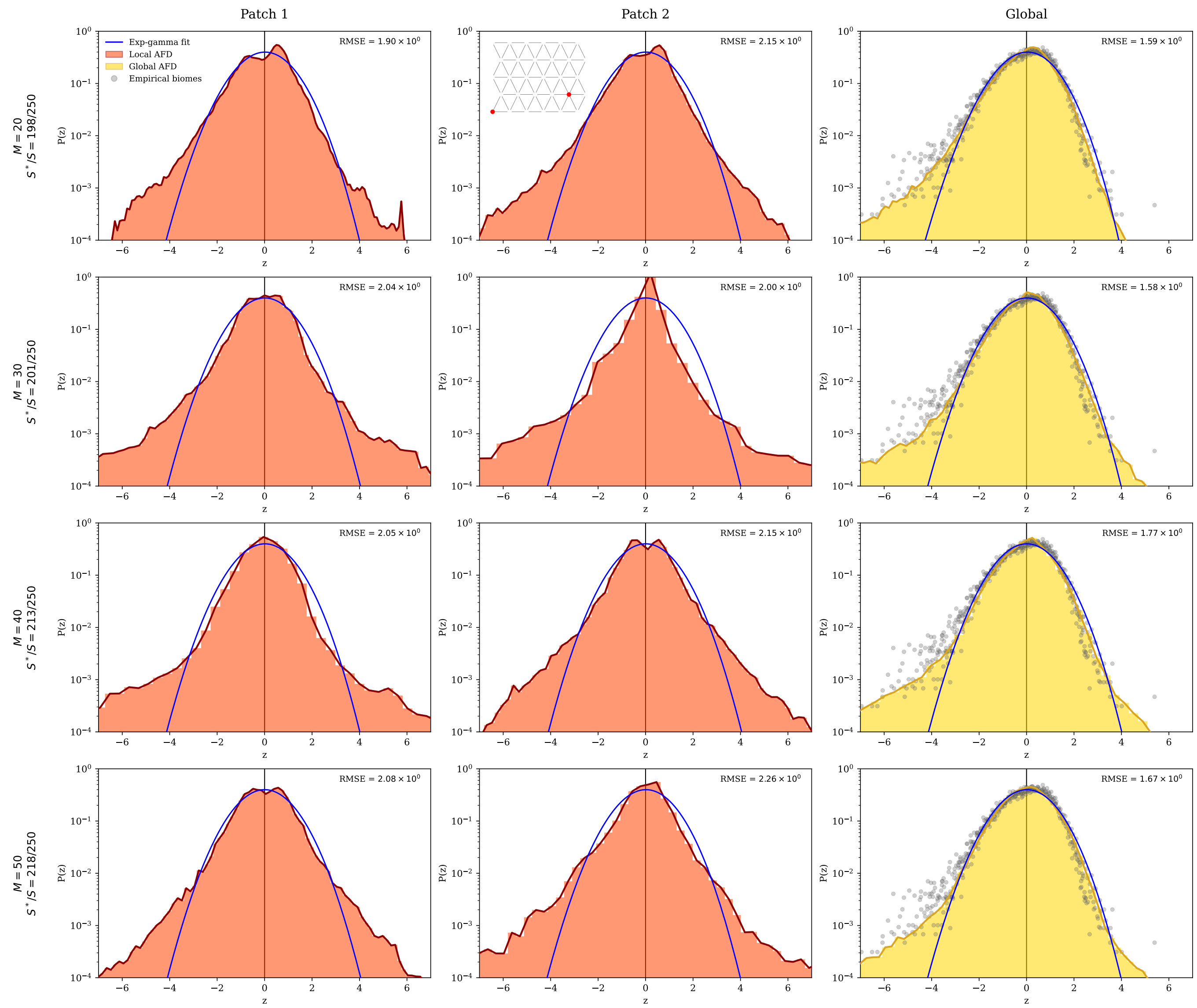


FIG. S18. **Abundance fluctuation distributions in a metacommunity with triangular-lattice embedding.** Same as Fig. S17, but with patches arranged on a triangular lattice. Patches are treated as vertices of the lattice, whose size $M$ varies across figure rows. Simulations are performed with $S = 250$, $D = 10^{-6}$, $\rho = 0.95$, $\mu S = 10$, $\sigma\sqrt{S} = 2$, and $\rho = 0.95$.

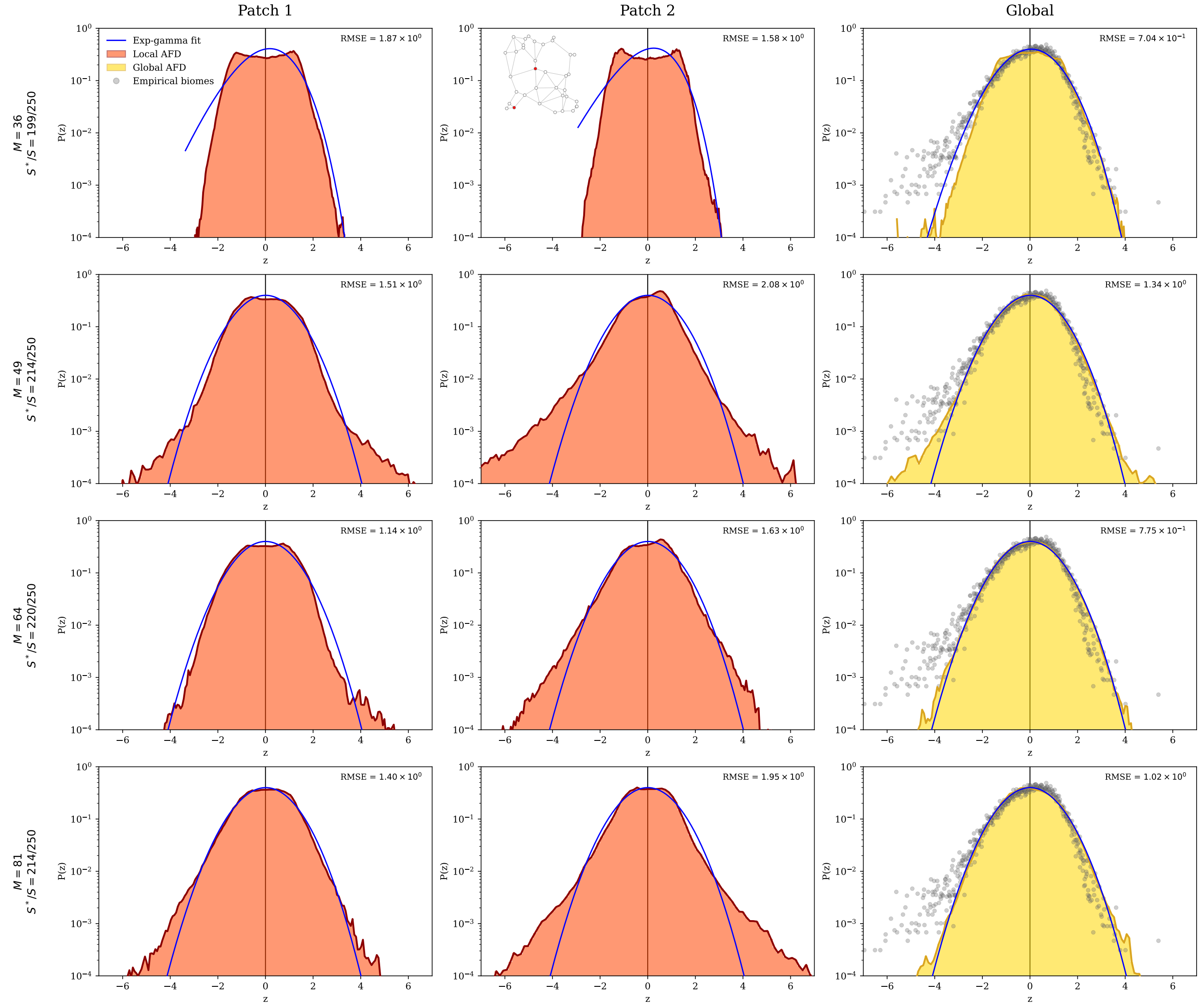


FIG. S19. **Abundance fluctuation distributions in the metacommunity model with a Gabriel graph.** Same as Figs. S17–S18, but with patches arranged on a Gabriel graph [4] following the metacommunity formulation and parametrisation introduced by O'Sullivan *et al.* [5], in which diffusion depends explicitly on the distance between patches. Patches are distributed uniformly within a square of side length $\sqrt{M}$ and connected whenever the disc having the segment between them as diameter contains no other patch. Diffusion coefficients are defined as $D_{uu} = -\lambda$, where $\lambda$ denotes the migration rate, while for distinct patches $(u \neq v)$ we set $D_{uv} = (\lambda/k_v)\exp(-d_{uv}/l)$, where $d_{uv}$ is the Euclidean distance between patches $u$ and $v$, $l = 0.5$ sets the characteristic migration length scale, and $k_v = \sum_{w\in\mathcal{N}(v)} \exp(-d_{vw}/l)$ is a normalisation constant summing over the set $\mathcal{N}(v)$ of Gabriel-graph neighbours of patch $v$, chosen so that the total outflow rate from any patch equals $\lambda$ regardless of its local connectivity. Simulations are performed with $\lambda = 0.01$, $S = 250$, $\rho = 0.95$, $\mu S = 10$, $\sigma\sqrt{S} = 2$, and the same extinction threshold used in the previous simulations.

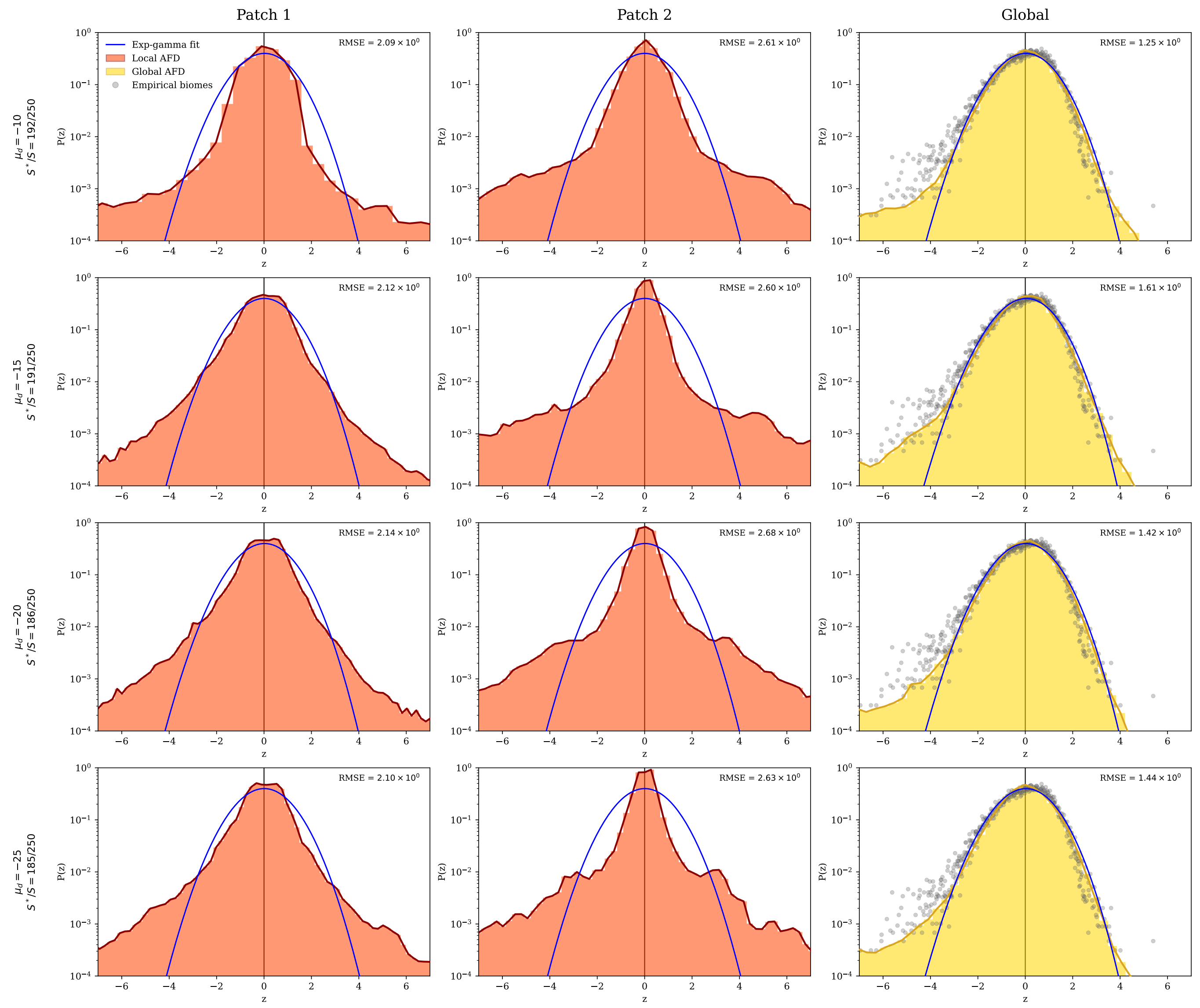


FIG. S20. **Abundance fluctuation distributions in the metacommunity model with a random diffusion matrix.** Same as Figs. S11–16, but with diffusion matrix entries $D^{uv}$ sampled from a lognormal distribution with mean $\mu_{\rm D} = 10$ and standard deviation $\sigma_{\rm D}$, which varies across rows. Simulations are performed with $S = 250$, $D = 10^{-6}$, $M = 25$, $\rho = 0.95$, $\mu S = 10$, and $\sigma\sqrt{S} = 2$.

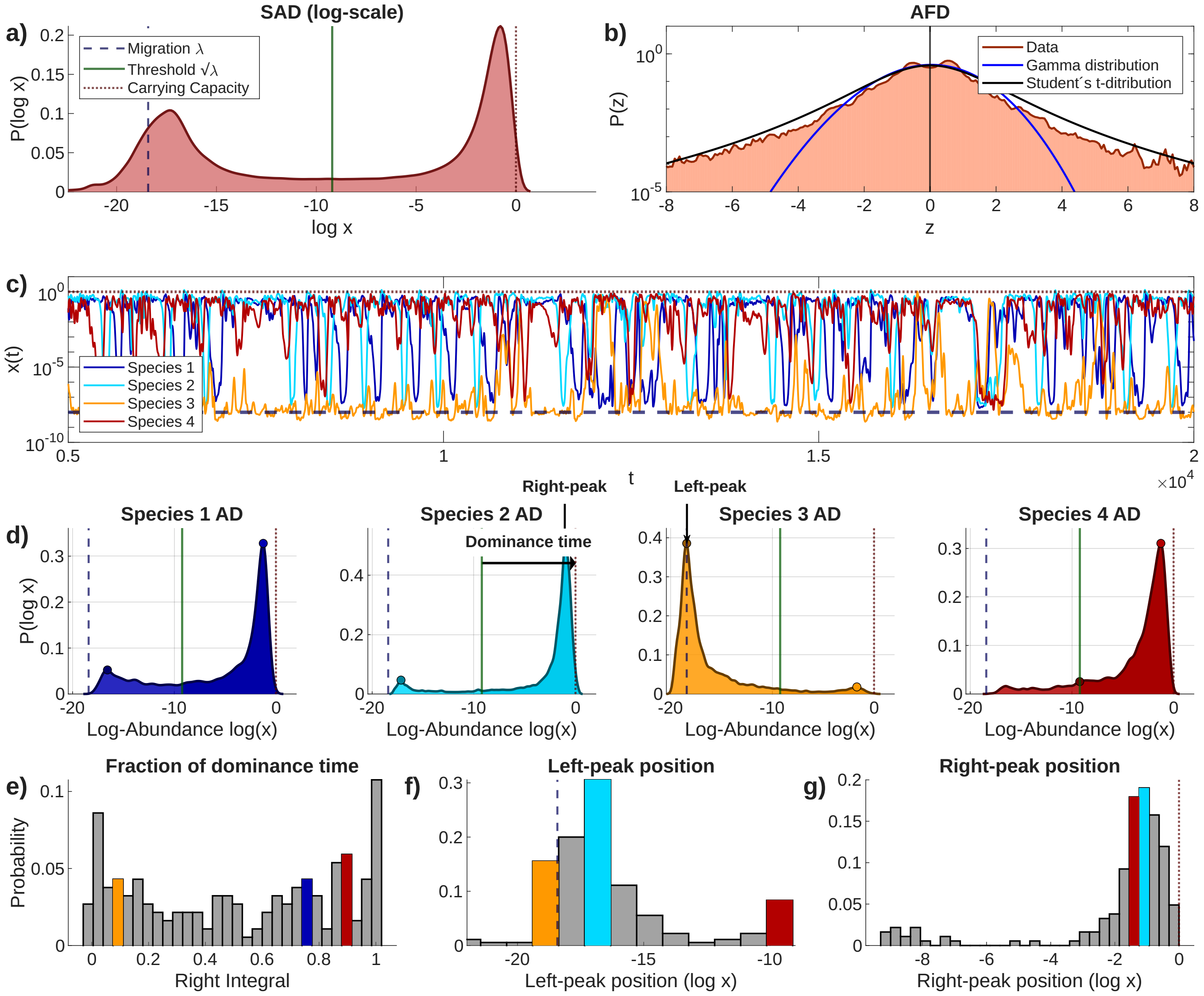


FIG. S21. **Macroecological characterization of the single patch dynamics in the metacommunity (a)** Species log-abundance distribution (distribution of log-abundances across time samples), aggregated over all species in the community. It displays a characteristic bimodal structure with an intermediate plateau. **(b)** Abundance fluctuation distribution (AFD), defined as the distribution of standardized log-abundances. Solid curves show fits to both the exp-gamma distribution and the Student's $t$ distribution. The exp-gamma fit performs markedly worse than the Student's $t$ fit. **(c)** Representative abundance time series for selected species. Abundances fluctuate between the migration floor $\lambda$ and the carrying capacity, displaying heterogeneous temporal behaviors ranging from small fluctuations to rapid turnover between rare and dominant states. Compared with Fig. S6, the lower value of $\sigma$ yields a more populated dominant community and a broader diversity of temporal behaviors. **(d)** Log-abundance distributions for the species shown in **(c)**. These distributions follow the general pattern shown in **(a)**, but the height and location of the peaks varies, reflecting a diversity of behaviors ranging from rare presence to high dominance. **(e)** Histogram of dominance times, defined as the fraction of time during which species abundances $x_i \geq \sqrt{\lambda}$. **(f), (g)** Distributions of the positions of the left and right peaks of the abundance distributions shown in **(d)**, respectively. Insets in **(d)** illustrate the definition of these quantities. Coloured bars indicate the locations in the histogram of the species illustrated in **(c)**, **(d)**. Results are obtained from simulations of Eq. (3) of the main text, with $S = 800$, $M = 10$, $\mu S = 10$, $\sigma\sqrt{S} = 2$, $x_{\mathrm{ext}} = 10^{-20}$, $D = 10^{-7}$ and $\rho = 0.95$.

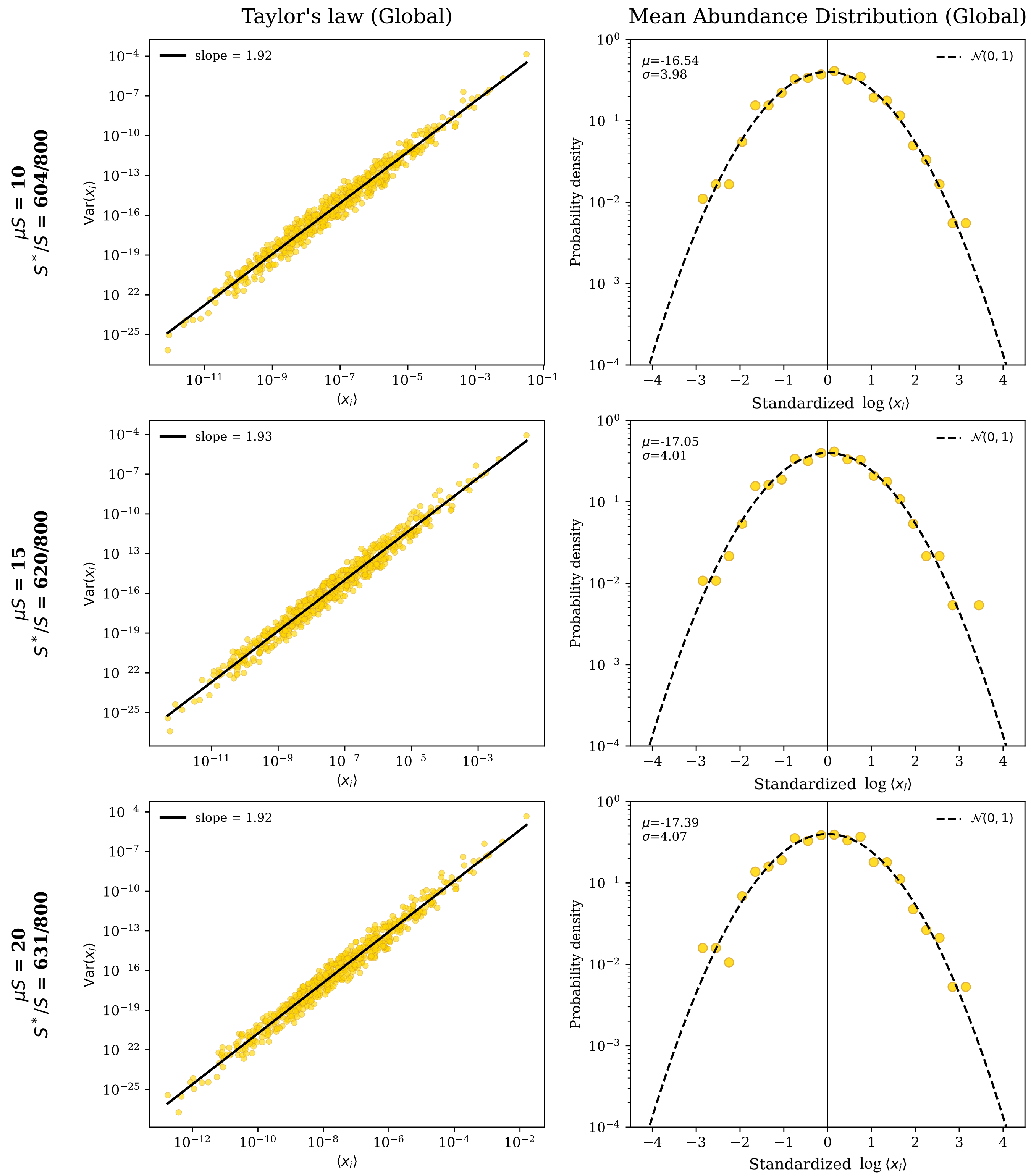


FIG. S22. **Taylor's law and the mean abundance distribution in the metacommunity model.** The metacommunity model reproduces the second and third macroecological laws reported by Grilli [6], namely Taylor's law with exponent close to 2 and a lognormal mean abundance distribution (MAD). Unlike the model with constant carrying capacities ($a_{ii} = 1$), reproducing these patterns requires heterogeneous carrying capacities. To this end, we assign species-specific carrying capacities drawn from a lognormal distribution, $K_i \sim \text{Lognormal}(K_\mu, K_\sigma^2)$, and shared across all patches. Interaction coefficients are then rescaled as $a_{ij} \to a_{ij}/K_j$, so that an isolated species reaches the equilibrium abundance $x_i^* = K_i$. **First column:** Taylor's law for the global (patch-aggregated) abundances, showing the expected power-law relationship between the temporal variance and mean abundance of each species, with exponent close to 2. **Second column:** Mean abundance distribution (MAD). We represent here the distribution across species of the logarithm of mean abundances along trajectories (standardized by subtracting their average and dividing by their standard deviation). The resulting distribution is well described by a Gaussian—meaning that the MAD is log-normally distributed. Rows correspond to different values of the mean interaction strength $\mu$. Simulations of Eq. (3) are performed with $S = 800$, $M = 10$, $\sigma = 2/\sqrt{S}$, $x_{\text{ext}} = 10^{-20}$, $D = 10^{-6}$, $\rho = 0.95$, $K_\mu = -14.0$, and $K_\sigma = 4.0$, where the last two parameters are taken from fits to empirical data reported in [6].

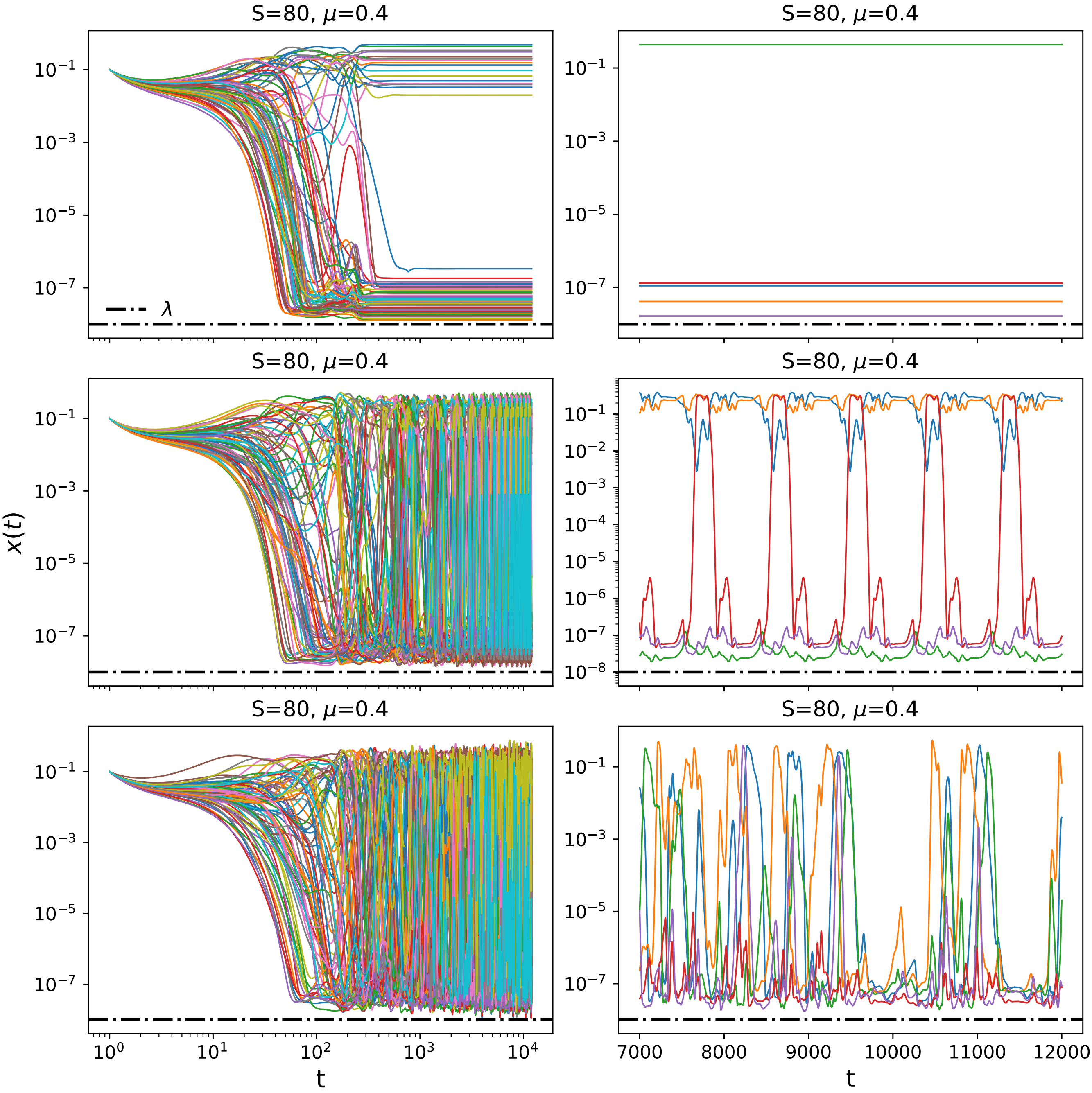


FIG. S23. **Dynamical phases of the open Lotka-Volterra model with constant migration.** The system displays an equilibrium phase (top row), as well as oscillatory behavior—including limit cycles (middle row) and chaotic fluctuations (bottom row)—within the multiple-attractor regime. Each panel shows simulated log-abundance dynamics as obtained from Eq. (2) of the main text. **Left:** Log-log plots of the full community dynamics, where the dashed black line indicates the migration level used in the simulations ($\lambda = 10^{-8}$), which sets a lower bound on species abundances. **Right:** Time series of five representative species, shown after discarding the initial transient, highlighting the richness and endogenous nature of the resulting fluctuations. Simulations are performed as described in Methods, with interaction coefficients sampled from a uniform distribution $a_{ij} \sim \mathcal{U}[0, 2\mu]$.

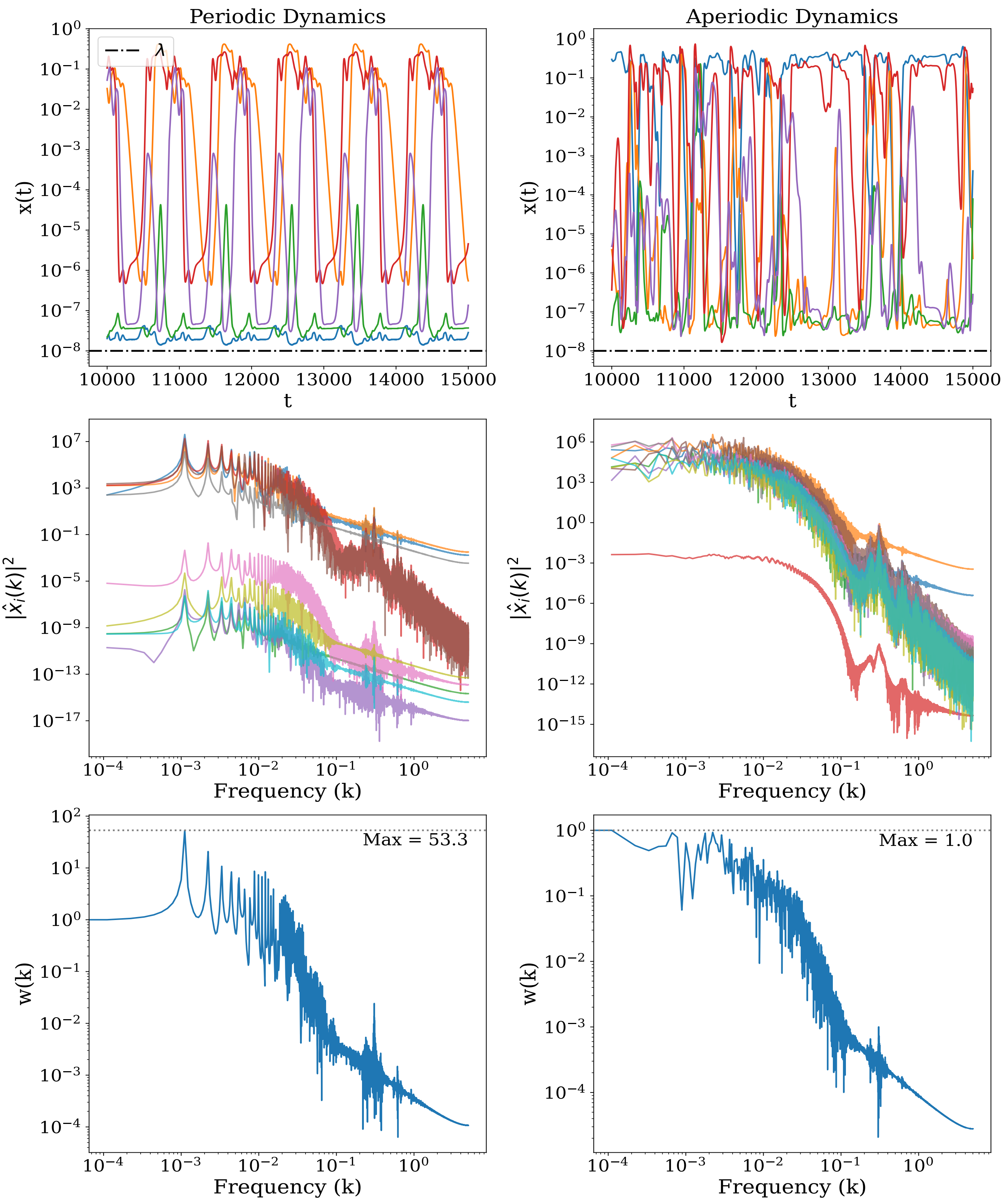


FIG. S24. **Periodicity analysis of the open gLV model with constant migration.** FFT-based analysis of periodic (left) and aperiodic (right) dynamics resulting from Eq. (2) of the main text, with $S = 80$, $\tau = 1$, $\lambda = 10^{-8}$, and interaction coefficients sampled from a uniform distribution $a_{ij} \sim \mathcal{U}[0, 2\mu]$ with $\mu = 0.4$. The first row shows the abundance time series for the two regimes, where different colours correspond to selected species and the black dot-dashed line indicates the migration level. The second row displays the squared modulus of the Fourier transform for 10 representative species, while the third row shows the quantity $w(k)$ (see Methods). A clear frequency peak is observed in the periodic regime, whereas it is absent in the aperiodic case.
